# Histone loaders CAF1 and HIRA restrict Epstein-Barr virus B-cell lytic reactivation

**DOI:** 10.1101/2020.04.28.067371

**Authors:** Yuchen Zhang, Chang Jiang, Stephen J. Trudeau, Yohei Narita, Bo Zhao, Mingxiang Teng, Rui Guo, Benjamin E Gewurz

## Abstract

Epstein-Barr virus (EBV) infects 95% of adults worldwide and causes infectious mononucleosis. EBV is associated with endemic Burkitt lymphoma, Hodgkin lymphoma, post-transplant lymphomas, nasopharyngeal and gastric carcinomas. In these cancers and in most infected B-cells, EBV maintains a state of latency, where nearly 80 lytic cycle antigens are epigenetically suppressed. To gain insights into host epigenetic factors necessary for EBV latency, we recently performed a human genome-wide CRISPR screen that identified the chromatin assembly factor CAF1 as a putative Burkitt latency maintenance factor. CAF1 loads histones H3 and H4 onto newly synthesized host DNA, though its roles in EBV genome chromatin assembly are uncharacterized. Here, we identified that CAF1 depletion triggered lytic reactivation and transforming virion secretion from Burkitt cells, despite strongly also inducing interferon stimulated genes. CAF1 perturbation diminished occupancy of histones 3.1, 3.3 and repressive H3K9me3 and H3K27me3 marks at multiple viral genome lytic cycle regulatory elements. Suggestive of an early role in establishment of latency, EBV strongly upregulated CAF1 expression in newly infected primary human B-cells prior to the first mitosis, and histone 3.1 and 3.3 were loaded on the EBV genome by this timepoint. Knockout of CAF1 subunit CHAF1B impaired establishment of latency in newly EBV-infected Burkitt cells. A non-redundant latency maintenance role was also identified for the DNA synthesis-independent histone 3.3 loader HIRA. Since EBV latency also requires histone chaperones ATRX and DAXX, EBV coopts multiple host histone pathways to maintain latency, and these are potential targets for lytic induction therapeutic approaches.

**IMPORTANCE:** Epstein-Barr virus (EBV) was discovered as the first human tumor virus in endemic Burkitt lymphoma, the most common childhood cancer in sub-Saharan Africa. In Burkitt lymphoma and in 200,000 EBV-associated cancers per year, epigenetic mechanisms maintain viral latency, where lytic cycle factors are silenced. This property complicated EBV’s discovery and facilitates tumor immunoevasion. DNA methylation and chromatin-based mechanisms contribute to lytic gene silencing. Here, we identify histone chaperones CAF1 and HIRA, which have key roles in host DNA replication-dependent and replication independent pathways, respectively, are each important for EBV latency. EBV strongly upregulates CAF1 in newly infected B-cells, where viral genomes acquire histone 3.1 and 3.3 variants prior to the first mitosis. Since histone chaperones ATRX and DAXX also function in maintenance of EBV latency, our results suggest that EBV coopts multiple histone pathways to reprogram viral genomes and highlights targets for lytic induction therapeutic strategies.

---

The gamma-herpesvirus Epstein-Barr virus (EBV) persistently infects nearly 95% of adults worldwide (1). EBV is the etiological agent of infectious mononucleosis and is also causally-associated with multiple human cancers, including endemic Burkitt lymphoma (eBL), Hodgkin lymphoma, post-transplant lymphoproliferative disease, HIV-associated lymphomas, nasopharyngeal carcinoma and gastric carcinoma (2). Tumor cells contain multiple copies of chromatinized, non-integrated, double-stranded DNA EBV genomes, where incompletely defined epigenetic pathways maintain a state of viral latency and in which most cells do not produce infectious virus.

EBV initiates lifelong infection by translocating across the tonsillar epithelium to colonize the B-cell compartment (3, 4). Virion deliver unchromatinized, encapsidated, linear EBV genomes to newly infected cells, which traffic to the nucleus. Upon nuclear entry, incoming genomes are circularized by host DNA ligases and chromatinized (1, 5, 6).

The EBV genome encodes nearly 80 proteins, most of which are highly immunogenic. To evade immune detection, EBV switches between latent and lytic genome programs, a hallmark of herpesvirus infection. Multiple layers of epigenetic regulation enable EBV to establish latency in newly-infected B-cells, in which a small number of viral encoded proteins and viral non-coding RNAs reprogram infected cell metabolism, growth and survival pathways (7–9). Within 3 days of infection, quiescent B-cells are reprogramed to become rapidly growing lymphoblasts that divide as frequently as every 8 hours (10–13).

According to the germinal center model(3), EBV-infected B-cells navigate the B-cell compartment to differentiate into memory cells, the reservoir for persistent EBV infection. To accomplish this, a series of EBV latency programs are used, in which combinations of Epstein-Barr nuclear antigens (EBNA), Latent Membrane Proteins (LMP) and ncRNA are expressed (1). Memory cells exhibit the latency I program, in which Epstein-Barr nuclear antigen 1 (EBNA1) is the only EBV-encoded protein expressed. EBNA1 tethers the EBV genome to host chromosomes and has key roles in propagation of viral genomes to daughter cells. EBNA1 is poorly immunogenic, facilitating immune escape of latency I cells.

Plasma cell differentiation is a trigger for EBV lytic reactivation. Induction of two viral immediate early gene transcription factors, BZLF1 and BRLF1, induce nearly 30 early genes important for production of lytic genomes (10, 14, 15). How these newly synthesized EBV genomes evade chromatinization by host histone loaders, including the heterotrimeric Chromatin Assembly Factor 1 (CAF1) complex that delivers newly synthesized H3/H4 dimers to host replication forks, is only partially understood (16, 17). EBV late genes are subsequently induced and include factors required for virion assembly and spread (10). Retrograde signals support ongoing lytic replication through subversion of chromatin-based repressors (18).

Most eBL cells utilize the latency I program, likewise enabling evasion of adaptive anti-EBV responses (19). Indeed, EBV was discovered as the first human tumor virus through eBL etiological studies, where the initial report noted that nearly all tumor cells did not produce infectious viral particles (20). With each S-phase, EBV genomes are copied once by host cell machinery and are then partitioned to daughter cells (21). Histone octamers consisting of two copies of histone 2A (H2A), H2B, H3 and H4 are loaded onto leading and lagging strands. CAF1 has key roles in loading histones onto newly replicated and damaged host DNA, whereas the histone chaperone HIRA is important for non-replicative histone loading onto host genomic sites (16, 17). Likewise, the chaperones alpha thalassemia/mental retardation syndrome X-linked chromatin remodeler (ATRX) and death domain-associated protein (DAXX) load histones onto telomeric and repetitive DNA. EBV tegument protein BNLF1 downmodulates ATRX/DAXX activity in newly infected cells (22), but ATRX/DAXX subsequently acquire important roles in the suppression of EBV lytic reactivation in latently infected cells (23).

Here, we characterize histone chaperone CAF1, HIRA, ATRX and DAXX roles in Burkitt EBV latency. We provide evidence that type I and II EBV strains co-opt each of these histone loaders to maintain latency via non-redundant roles. EBV upregulated each of the three CAF1 subunits in newly infected primary human B-cells, and CAF1 was found to have key roles in establishment of latency in a Burkitt EBV infection model. Chromatin immunoprecipitation assays support key CAF1 roles in deposition of repressive histone marks on EBV genome lytic control elements. These data further support key chromatin roles in regulation of the EBV lytic switch.

## RESULTS

### The Histone Loader CAF1 is important for Burkitt lymphoma EBV latency maintenance

To gain insights into host factors important for the maintenance of EBV latency, we recently performed a human genome wide CRISPR screen (24). Briefly, Cas9+ EBV+ Burkitt P3HR-1 cells were transduced with the Avana single guide RNA (sgRNA) library, which contains four independent sgRNAs against nearly all human genes. Cells with de-repressed plasma membrane (PM) expression of the EBV late lytic antigen gp350, indicative of latency reversal, were sorted at Days 6 and 9 post-transduction. sgRNAs significantly enriched in the sorted versus input cell population were identified. The STARS algorithm identified 85 statistically significant hits at a p<0.05 and fold change > 1.5 cutoff (Fig. 1A)(24, 25). Unexpectedly, genes encoding two subunits of the histone loader CAF1 complex were amongst top screen hits (Fig. 1A-C).

**Figure 1.**
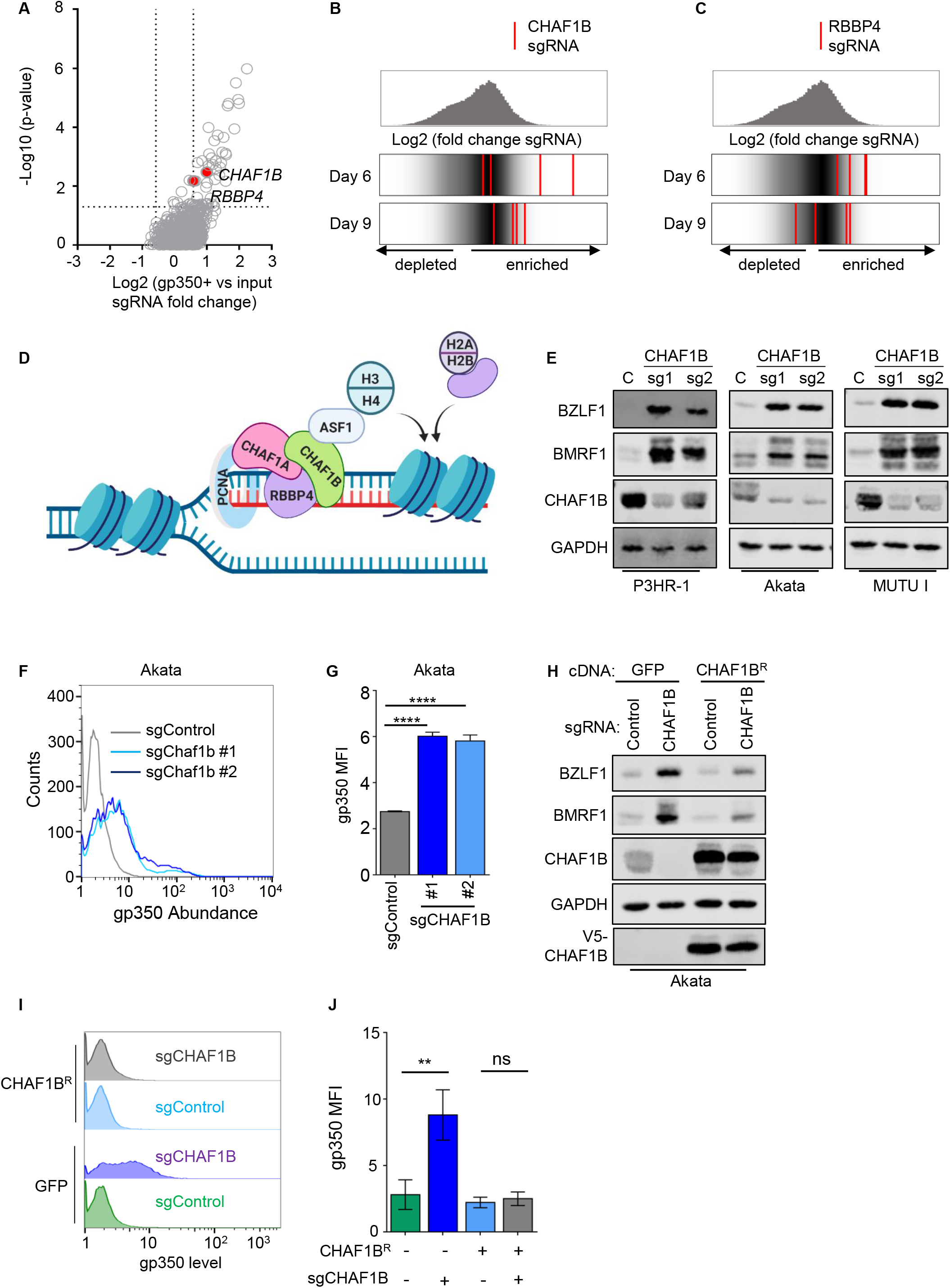
CHAF1B depletion triggers EBV lytic gene expression in Burkitt cells. (A) Volcano plots of CRISPR screen(24) −Log10 (p-value) and Log2 (fold-change of gp350+ vs input library sgRNA abundance) on Day 6 post Avana library transducton. CAF1 subunits. (B, C) Top: Distribution of Log2 (fold-change gp350+ versus input library sgRNA abundance) at Day 6 (B) or Day 9 (C) post sgRNA expression. Bottom: Log2 fold change for the four CHAF1B (B) or RBBP4 (C) targeting sgRNAs (red lines), overlaid on gray gradient depicting overall sgRNA distributions at CRISPR screen Days 6 versus 9. Average values from two screen biological replicates are shown. (D) Model of DNA replication-dependent histone H3 and H4 loading by CAF1 and ASF1. Also shown are the CAF1 binding partner PCNA clamp and a histone chaperone loading histones H2A/H2B onto DNA. (E) Immunoblot analysis of whole cell lysates (WCL) from P3HR-1, Akata and MUTU I Burkitt cells expressing control or CHAF1B sgRNAs. (F) FACS analysis of plasma membrane (PM) gp350 expression in Akata cells expressing control or CHAF1B sgRNAs. (G) Mean ± standard deviation (SD) PM gp350 mean fluorescence intensities (MFI) from n=3 replicates, as in (F). **** p < 0.0001. (H) Immunoblot analysis of WCL from Akata cells expressing GFP or V5-epitope tagged CHAF1B rescue cDNA (CHAF1B^R^) and the indicated sgRNAs. (I) FACS analysis of PM gp350 expression in Akata cells that stably express GFP or CHAF1B^R^ and the indicated sgRNAs. (J) Mean ± SD PM gp350 MFI values from n=3 replicates, as in (I). Cells expressed GFP where not indicated to express CHAF1B^R^, and cells expressed sgControl where not indicated to express sgCHAF1B. ** p < 0.01, ns, not significant. Blots in E and H are representative of n=3 replicates.

The heterotrimeric CAF1 complex, comprised of CHAF1A, CHAF1B and RBBP4 subunits, delivers histone H3/H4 dimers to the replication fork during cell cycle S-phase, typically together with histone chaperone ASF1a (17, 26) (Fig. 1D). CAF1 has well-established roles in maintenance of heterochromatin and cell identity, but its function in regulating EBV latency has not yet been investigated. Therefore, it was notable that multiple sgRNAs targeting *CHAF1B* and *RBBP4* were enriched amongst gp350+ sorted cells at Days 6 and 9 post-Avana library transduction (Figs. 1B-C). A sgRNA targeting *CHAF1A* was also enriched in gp350+ cells at the Day 6 timepoint (Fig. S1A). The identification of multiple sgRNAs targeting CAF1 subunits suggested an important CAF1 role in maintenance of EBV latency. Notably, Burkitt lymphoma are the fastest growing human tumor, and newly synthesized EBV genomes must be reprogrammed to maintain latency I with each cell cycle.

To validate screen hit CAF1 roles in the maintenance of BL EBV latency, control or independent *CHAF1B* targeting sgRNAs were expressed in P3HR-1, Akata and MUTU I Cas9+ tumor-derived endemic Burkitt lymphoma cell lines. In each of these, CHAF1B depletion induced immediate early BZLF1 and early BMRF1 expression (Fig. 1E). CHAF1B depletion significantly induced all seven EBV lytic transcripts measured by qRT-PCR (Fig. S1B). Since Akata and MUTU I harbor type I EBV, whereas P3HR-1 carries type II EBV, these data suggest a conserved CAF1 roles in maintenance of EBV latency. Likewise, CHAF1B depletion induced gp350 plasma membrane expression on most Akata cells examined by flow cytometry (Fig. 1F-G), suggesting that Burkitt cell CAF1 loss triggers a full EBV lytic cycle. CHAF1B depletion also induced gp350 expression on MUTU I cells (Fig. S1C-D).

We next validated on-target CRISPR effects through a cDNA rescue approach. A point mutation was engineered into the *CHAF1B* cDNA proto-spacer adjacent motif (PAM) site targeted by sgRNA #1 to abrogate Cas9 editing. Akata cells with stable control GFP vs V5-epitope tagged CHAF1B rescue cDNA (CHAF1B^R^) were established. Effects of control vs *CHAF1B* targeting sgRNA were tested. Interestingly, depletion of endogenous CHAF1B de-repressed BZLF1, BMRF1 and gp350 in control cells, but failed to do so in cells with CHAF1B^R^ rescue cDNA expression (Figs. 1H-J). Similar cDNA rescue results on BZLF1 and BMRF1 expression were evident in MUTU I cells (Fig. S1E). These results suggest that CHAF1B is necessary for EBV latency in Burkitt cells, perhaps in loading histone H3/H4 onto newly synthesized episomes.

### CHAF1B Perturbation Induces EBV Genome Lytic Replication and IFN Stimulated Genes

EBV lytic replication is controlled on many levels and partial lytic cycle induction is often observed. Therefore, we next examined whether CAF1 perturbation was sufficient to induce a productive lytic replication cycle. RNAseq was performed on Akata cells at Day 6 post-sgRNA expression, and demonstrated significant induction of EBV 77 lytic cycle genes (Fig. 2A). Consistent with induction of a full lytic cycle, CHAF1B depletion induced intracellular EBV genome amplification, albeit to a level less than observed with Akata immunoglobulin (Ig) crosslinking. Likewise, CHAF1B sgRNAs induced secretion of DNAse-resistant EBV genomes, demonstrating encapsidation (Fig. 2B). Similar results were observed in MUTU I and P3HR-1 cells, suggesting conserved CAF1 roles in type I and II EBV latency regulation (Fig. S2A-B). In support of on-target CRISPR effects, expression of the PAM site mutant CHAF1B cDNA rescue construct prevented EBV genome copy number increase with editing of endogenous *CHAF1B* (Fig. S2C). Furthermore, addition of supernatant from CHAF1B depleted, but not control Akata cells, stimulated human B-cell aggregation and growth transformation (Fig. 2C).

**Figure 2.**
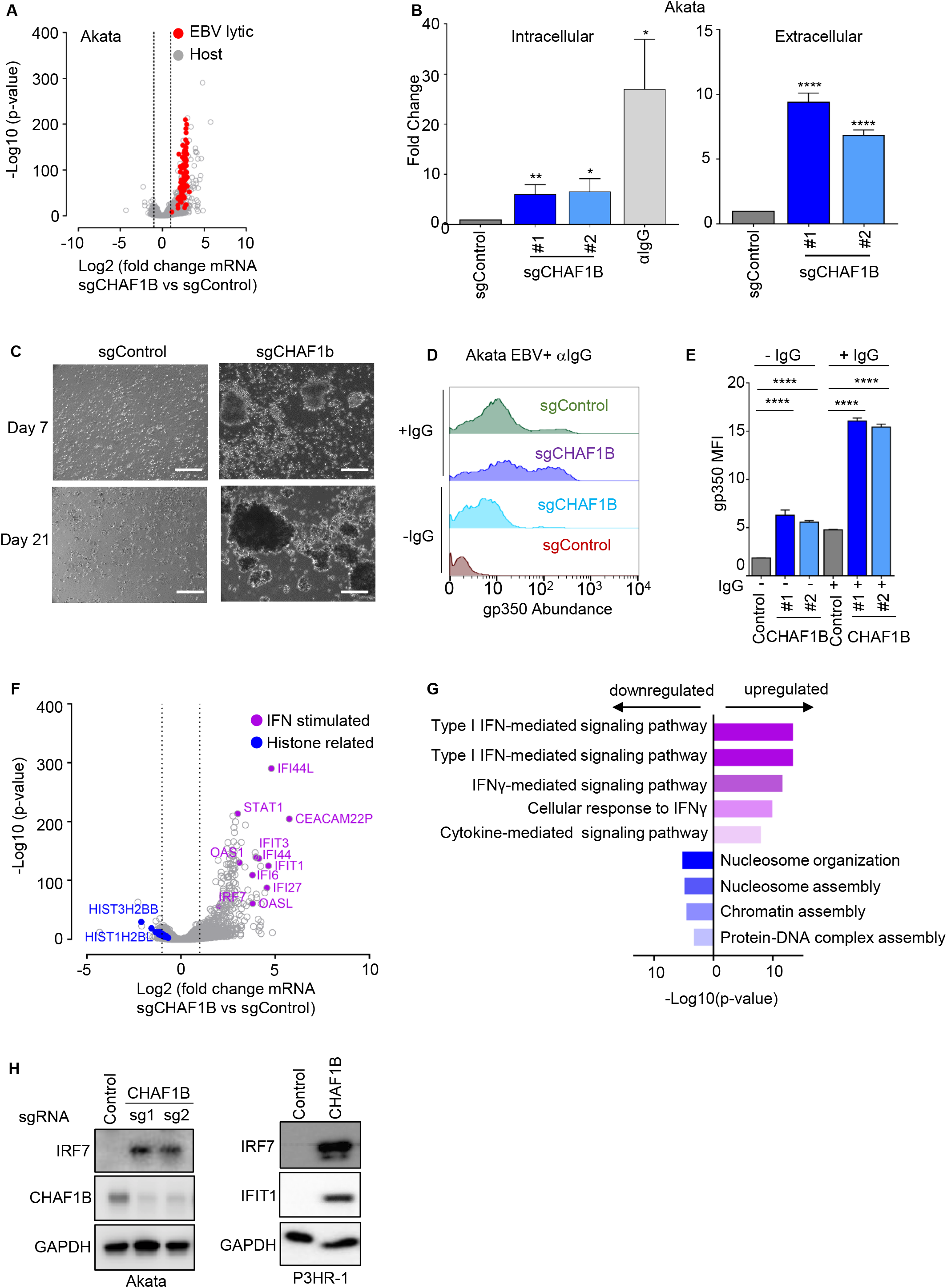
CHAF1B depletion triggers Burkitt cell EBV lytic reactivation and interferon stimulated gene expression. (A) Volcano plot comparing RNAseq −Log10 (p-value) versus Log2 (fold-change sgCHAF1B vs sgControl mRNA abundance) from n=3 replicates. Significantly changed EBV lytic gene values are shown in red, host genes are shown in gray. (B) qPCR analysis of EBV intracellular or DNase-treated extracellular genome copy number from Akata cells expressing control or CHAF1B sgRNAs. Total genomic DNA was extracted at Day 6 post lentivirus transduction or 48h post stimulation by anti-IgG (10μg/ml). Mean ± SD values from n=3 biologically independent replicates are shown. *p<0.05, **p<0.01, ****p<0.0001. (C) Phase microscopy images of human primary B cells at Day 7 or 21 post-inoculation with cell culture supernatant from Akata cells expressing control or CHAF1B sgRNAs. White scale bar=100μm. (D) Representative FACS plots of PM gp350 expression in Akata cells expressing control or CHAF1B sgRNAs and in the absence or presence of αIgG (10μg/ml) for 48 hours. (E) Mean ± SD PM gp350 MFI values from n=3 replicates of Akata with indicated sgRNAs and αIgG stimulation, as in (D). **** p < 0.0001. (F) Volcano plot comparing RNAseq −Log10 (p-value) versus Log2 (fold-change sgCHAF1B vs sgControl mRNA abundance) from n=3 replicates. Purple circles indicate selected interferon (IFN) stimulated genes and blue circles indicate histone related genes. (G) Enrichr pathway analysis of gene sets significantly upregulated (purple bars) or downregulated (blue bars) by CHAF1B sgRNA expression. Shown are the −Log10 (p-values) from Enrich analysis of triplicate RNAseq datasets, using Fisher exact test. See also Table S1. (H) Immunoblot analysis of WCL from Akata or P3HR-1 cells expressing control or CHAF1B sgRNAs, for the IFN stimulated genes IFIT1 and IRF7, CHAF1B or GAPDH, as indicated.ulated gene expression. Blots in H are representative of n=2 replicates.

RNAseq analysis also demonstrated robust up-regulation of EBV latency III transcripts in response to CHAF1B depletion. EBNA1, 2, 3A, 3B, 3C, LMP1, LMP2A and LMP2B were each significantly up-regulated (Fig. S3A and Table S1). While these transcripts are upregulated by EBV lytic reactivation (27), the magnitude of mRNA upregulation suggests that CAF1 may also have important roles in chromatin-based silencing of the latency III program.

To test whether CAF1 perturbation and Ig-crosslinking synergistically induce lytic replication, control versus CHAF1B sgRNAs were expressed in Cas9+ Akata cells in the absence or presence of αIgG. Interestingly, Ig-crosslinking induced higher levels of PM gp350 and intracellular/extracellular EBV genome copy numbers in cells depleted for CHAF1B than in control cells (Fig. 2D-E and S2D). Similar results were obtained with IgM cross-linking in MUTU I cells (Fig. S2E-F). These results suggest that CAF1 not only maintains EBV latency in unstimulated cells, but also limits the extent of lytic reactivation in upon B-cell receptor activation.

We next examined changes in host mRNAs following CHAF1B depletion. Interestingly, multiple interferon stimulated genes (ISGs) were amongst the most highly CHAF1B sgRNA induced host genes, including IFIT1, IFIT3, IFI44, IFI44L, IRF7 and STAT1, and GO analysis identified Type I interferon-mediated signaling pathway as the most highly upregulated pathway (Fig. 2F-G, S3B). By contrast, mRNAs encoding histones and histone-related genes were amongst the most strongly downmodulated by CHAF1B depletion (Fig. 2F-G, S3C), perhaps as a result of a negative feedback in response to diminished CAF1 activity. CHAF1B-mediated upregulation of IRF7 and IFIT1 was validated at the protein level (Fig. 2H). Interestingly, ISG upregulation was not observed at the mRNA or protein level in Akata cells upon immunoglobulin-crosslinking induced EBV lytic reactivation (28–30). These results suggest that that EBV lytic replication itself does not underlie this host response, at least when triggered by Ig-crosslinking.

### Depletion of CAF1 Subunits CHAF1A and RBBP4 triggers EBV lytic replication

CHAF1B assembles together with CHAF1B and RBBP4 subunits with 1:1:1 stoichiometry (31). CHAF1A targets CAF1 to the replication fork through interaction with proliferating cell nuclear antigen (PCNA), associates with histone deacetylases, and has roles in DNA repair and in heterochromatin maintenance (17, 32). While RBBP4 has been implicated in CAF1 activity (33), it also has additional epigenetic roles, including within the NURD transcriptional repressor complex(34).

To investigate whether the CAF1 subunits CHAF1A and RBBP4 were similarly important for the maintenance of Burkitt EBV latency I, we tested the effects of the top two Avana library sgRNAs targeting the genes encoding each. Depletion of RBBP4 or CHAF1A by either sgRNA induced all seven EBV lytic genes surveyed by qPCR (Fig. S4A-B) and induced BZLF1, BMRF1 and gp350 at the protein level (Fig. 3A-F). RBBP4 or CHAF1A depletion likewise de-repressed EBV lytic gene expression in P3HR-1 and MUTU I (Fig. S4C-D). To determine effects of RBBP4 or CHAF1A depletion on EBV genome amplification, viral load analysis was performed. RBBP4 and CHAF1A sgRNAs significantly increased intracellular and DNAse-treated extracellular EBV genome copy numbers in three Burkitt cell lines (Fig. 3G-H, S4E-F). Taken together, these results suggest that all three CAF1 subunits are critical for EBV latency in Burkitt lymphoma cells, perhaps acting to re-program newly synthesized EBV episomes with each cell cycle.

**Figure 3.**
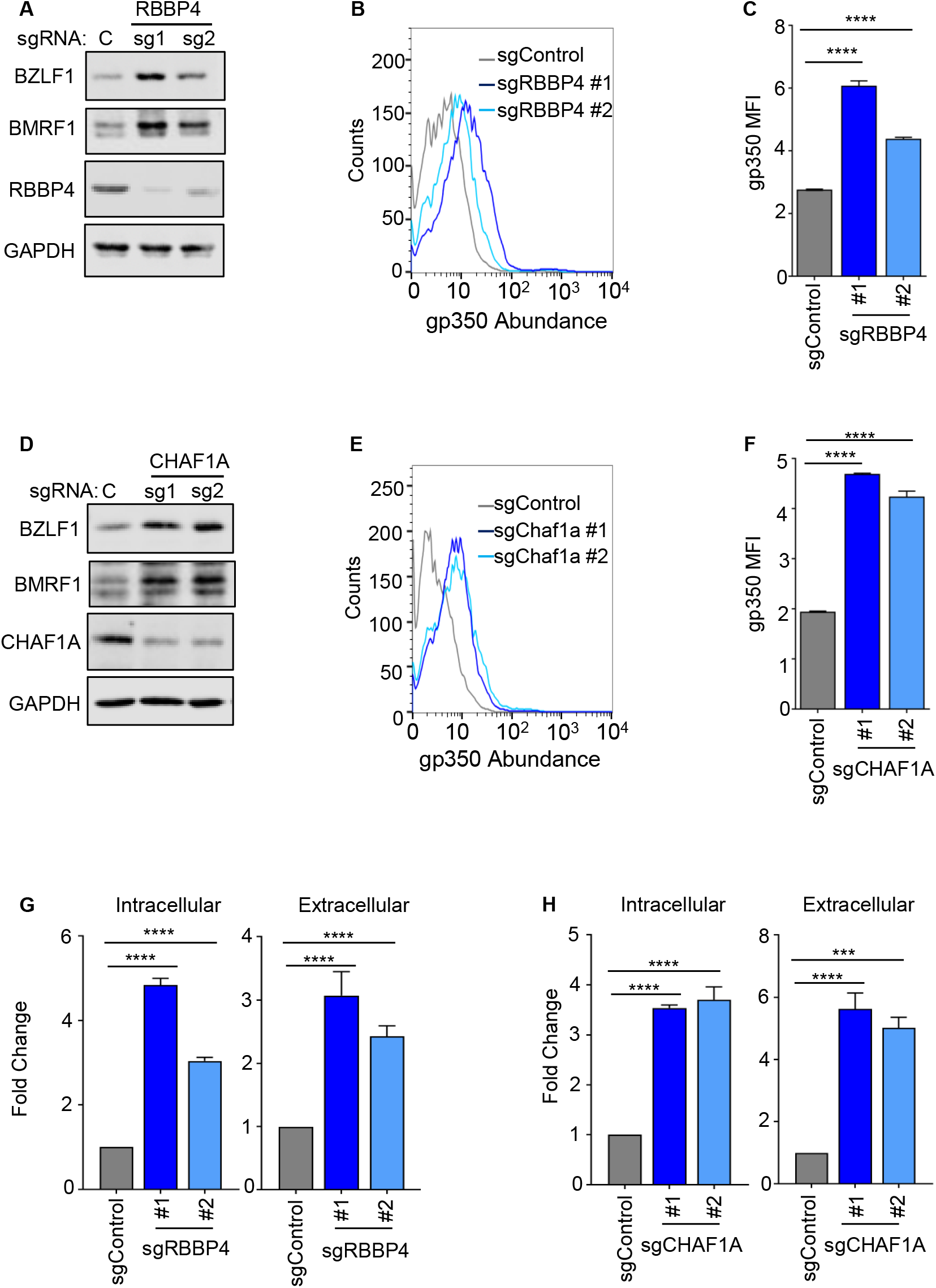
CAF1 subunits RBBP4 and CHAF1A are necessary for Burkitt cell EBV latency. (A) Immunoblot analysis of WCL from Akata cells expressing control or RBBP4 sgRNAs. (B) FACS analysis of PM gp350 expression in Akata cells expressing control or RBBP4 sgRNAs. (C) Mean ± SD PM gp350 MFI values from n=3 replicates of Akata with the indicated sgRNAs, as in (B). **** p < 0.0001. (D) Immunoblot analysis of WCL from Akata expressing control or CHAF1A sgRNAs. (E) FACS analysis of PM gp350 expression in Akata cells expressing control or CHAF1A sgRNAs. (F) Mean ± SD PM gp350 MFI values from n=3 replicates of Akata with indicated sgRNAs, as in (E). **** p < 0.0001. (G and H) qPCR analysis of EBV intracellular or DNAse-treated extracellular genome copy number from Akata expressing control, RBBP4 (G) or CHAF1A (H) sgRNAs. Total genomic DNA was extracted at Day 6 post lentivirus transduction. Mean ± SD values from n=3 replicates are shown. ****p<0.0001. Blots in A and D are representative of n=3 replicates.

### Roles of EBV-induced CAF1 in establishment of B-cell latency

Since CAF1 has key histone deposition roles in the contexts of DNA replication or repair, we asked whether CAF1 subunits are expressed in resting or in newly-infected primary human B-cells. Using data from recently published RNAseq and proteomic maps of EBV-mediated primary B-cell growth transformation (35, 36), we noticed that there was little expression of CHAF1A or CHAF1B in the resting B-cells, but that each are upregulated by 2 days post-infection (Fig. 4A, S5A). RBBP4 appears to have a higher basal level, perhaps reflective of its additional epigenetic roles beyond CAF1, but is also EBV-upregulated (Fig. S5A). Immunoblot analysis demonstrated strong CHAF1B upregulation between 2 and 4 days post-infection (Fig. 4B), at which point newly infected cells begin to rapidly proliferate as they transition from the EBV pre-latency program to latency IIb (12, 37). Published LCL Chip-seq data (38–41) showed Epstein-Barr nuclear antigens 2, LP, 3A, 3C and LMP1-activated NF-kB subunit occupancy at or near the *CHAF1A, CHAF1B* and *RBBP4* promoters (Fig. 4C, S5B-C). MYC occupancy was also notable at the *CHAF1A* and *RBBP4* promoters in Burkitt-like P493 B-cells (42). We therefore speculate that these EBNAs, which are expressed in the EBV pre-latency and latency IIb programs, have important roles in EBV-mediated CAF1 upregulation in newly infected primary B-cells.

**Figure 4.**
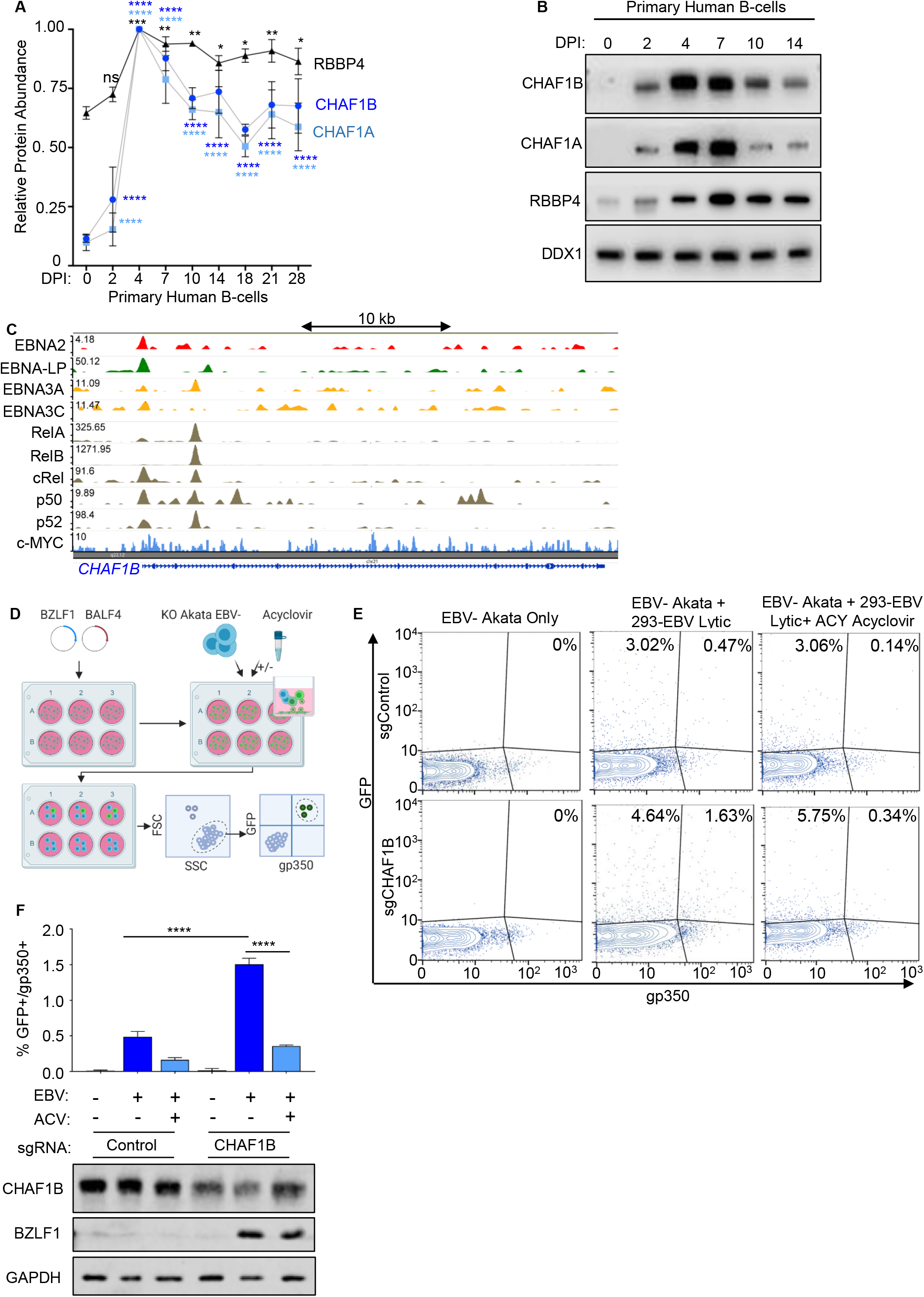
CAF1 complex restricts lytic cycle after EBV infection in primary human B cells. (A) CHAF1B, CHAF1A and RBBP4 relative protein abundances detected by tandem-mass-tag-based proteomic analysis of primary human B-cells at rest and at nine time points after EBV B95.8 infection at a multiplicity of infection of 0.1proteomic analysis at rest and at nine time points after EBV B95.8 strain infection of primary human peripheral blood B-cells at a multiplicity of infection of 0.1. Data represent the average +/- SEM for n=3 independent replicates(35). For each protein, the maximum level detected across the time course was set to a value of 1. (B) Immunoblot analysis of WCL from primary B cells infected with B95.8 EBV at Days 0, 2, 4, 7, 10 and 14 post-infection. (C) GM12878 ChIP-seq signals of EBV-encoded EBNA2, EBNA-LP, EBNA3A, EBNA3C, LMP1 activated RelA, RelB, cRel, p50, p52 NF-kB subunits or c-Myc at the *CHAF1B* locus. Track heights are indicated in the upper left. (D) Schematic diagram of cell co-culture system for newly-infected Burkitt cell EBV latency establishment. Transfection of *BZLF1* and *BALF1* expression vectors triggers lytic reactivation in EBV+ 293 cells with a recombinant viral genome that harbors a GFP marker. EBV-uninfected (EBV-) Akata cells are then co-cultured with induced 293 cells, in the absence or presence of acyclovir. 48 hours later, cells are analyzed by FACS for expression of GFP and the late lytic antigen gp350, which is expressed in cells that fail to establish latency. (E) Control or CHAF1B KO Akata EBV-cells were co-cultured with HEK-293 2-8-15 cells harboring GFP-EBV. Cells were mock treated or treated with 50μg/ml of acyclovir. Cells were then subjected to GFP and PM gp350 FACS. FSC and SSC parameters were used to gate out the contaminated HEK293 cells from Akata EBV negative cells. GFP vs gp350 dot plots from a representative replicate were shown. (F) Top: Mean ± SD PM gp350 MFI values from n=3 replicates of co-cultured Akata EBV-cells with the indicated experimental conditions, as in (D). **** p < 0.0001. Bottom: Immunoblot analysis of WCL from Akata EBV-cells co-cultured 293 cells under the indicated experimental conditions.

We next tested CAF1’s roles in the establishment of EBV latency. Since it is not currently possibly to do CRISPR editing in resting primary B-cells, we instead used an EBV-negative (EBV-) subclone of Akata Burkitt cells, which were established during serial passage of the original EBV+ Akata tumor cells (43). It has previously been shown that latency I is established upon re-infection of these cells by EBV *in vitro* (44). However, since EBV-Akata cells are difficult to infect with purified EBV, we developed a co-culture system to increase infection efficiency (Fig 4D). EBV-Akata were co-cultured with EBV+ 293 producer cells, which carry a recombinant EBV bacterial artificial chromosome (BAC) system that includes a GFP marker (45). Lytic replication was induced in a monolayer of adherent 293-EBV+ cells by transfection of genes encoding BZLF1 and BALF4. Induced 293 cells were then co-cultured with Akata EBV-cells 24 hours post-transfection. EBV infection frequency was monitored by FACS 48 hours later, using GFP as a readout, and PM gp350 positivity was used as a marker for cells with lytic replication.

Using this co-culture system, ~3.5% of control Akata cells were infected, as judged by expression of the GFP marker, and 0.47% of cells were positive for the gp350 lytic antigen. By comparison, ~6% of CHAF1B depleted cells were infected and 1.63% had gp350 PM expression (Fig. 4E). Most gp350 expression was suppressed by addition of acyclovir to the co-culture system, suggesting it was expressed as a late lytic gene rather than delivered by incoming or attached EBV (Fig. 4E). Further suggesting an important CAF1 role in establishment of latency in Akata cells, BZFL1 was more highly expressed in CHAF1B-depleted than control cells. As expected, expression of this immediate early gene was not blocked by acyclovir (Fig. 4F). These data are consistent with a model in which CAF1 has key roles in reprogramming the epigenetic state of newly infected B-cells. However, it is possible that CAF1 plays an earlier role in this rapidly growing Akata system than in primary B-cells, where the first mitosis occurs 72 hours post-infection.

### The Histone Chaperone HIRA exerts non-redundant Burkitt cell maintenance of EBV Latency roles

The histone loader Histone Regulatory Homologue A (HIRA) interacts with ASF1a and preferentially loads histone H3.3/H4 complexes onto DNA in a replication-independent manner throughout the cell cycle, for example at areas of active transcription (17, 46). HIRA regulates the alpha-herpesvirus herpes simplex virus and the beta-herpesvirus cytomegalovirus latency (47–50). HIRA is also implicated in maintenance of HIV latency (51), but to our knowledge has not been investigated in the regulation of gamma-herpesvirus latency.

To explore potential HIRA roles in the maintenance of EBV latency, we tested effects of HIRA depletion in P3HR-1, Akata and MUTU I cells. In each of these BL, CRISPR *HIRA* editing by either of two Avana sgRNAs rapidly upregulated BZLF1 and BMRF1 in Akata, P3HR-1 and MUTU I cells (Fig. 5A). HIRA depletion also upregulated PM gp350 abundance, albeit to a lesser extent than observed with CAF1 perturbation, perhaps explaining why our CRISPR screen was more sensitive to CAF1 perturbation (Fig. 5B-C). HIRA sgRNAs increased expression of all seven EBV lytic mRNAs quantified by qPCR (Fig. S6A) as well as EBV genome copy number (Fig. 5D). Supernatants from HIRA-depleted cells induced primary human B-cell clumping, though clusters were generally smaller than observed with CHAF1B KO, likely reflecting lower titer of secreted EBV (Fig. 5D). Taken together, these results indicated that HIRA and CHAF1B have non-redundant roles in the maintenance of Burkitt EBV latency, as depletion of either triggers lytic reactivation.

**Figure 5.**
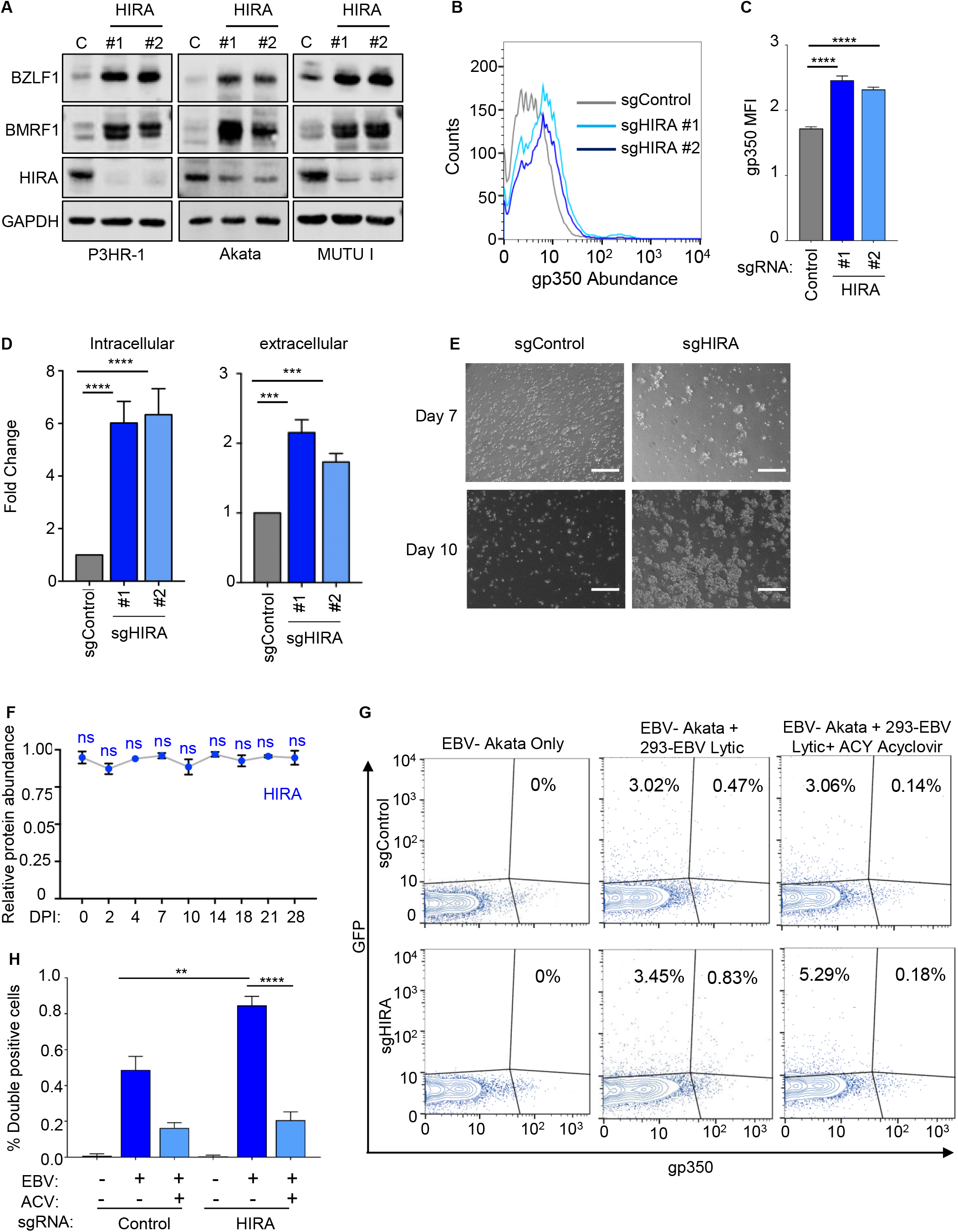
Histone 3.3 chaperone HIRA restricts Burkitt EBV lytic reactivation. (A) Immunoblot analysis of WCL from P3HR-1, Akata EBV+ or MUTU I BL cells expressing control or HIRA sgRNAs. (B) FACS analysis of PM gp350 expression in Akata EBV positive cells expressing control or HIRA sgRNAs. (C) Mean ± SD PM gp350 MFI values from n=3 replicates of Akata with indicated sgRNAs, as in (B). **** p < 0.0001. (D) qPCR analysis of EBV intracellular or DNAse-resistant extracellular genome copy number from Akata EBV+ cells expressing control or HIRA sgRNAs. Total genomic DNA was extracted at Day 6 post lentivirus transduction. Mean ±SD values from n=3 replicates are shown. ****p<0.0001. (E) Phase microscopy images of human primary B cells at Day 7 or 10 post-inoculation with cell culture supernatant from Akata cells expressing control or HIRA sgRNAs. White scale bar=100μm. (F) HIRA relative protein abundances detected by tandem-mass-tag-based proteomic analysis of primary human B-cells at rest and at nine time points after EBV B95.8 infection at a multiplicity of infection of 0.1. Data represent the average +/- SEM for n=3 independent replicates(35). For each protein, the maximum level detected across the time course was set to a value of 1. (G) Control or HIRA KO Akata EBV-cells were co-cultured with HEK-293 2-8-15 cells harboring recombinant EBV encoding a GFP marker. Cells were mock-treated or treated with 50μg/ml of acyclovir. Cells were then subjected to FACS for GFP or PM gp350. GFP vs gp350 dot plots from a representative replicate were shown. (H) Mean + SD PM gp350 MFI values from n=3 replicates Akata EBV-cells of co-cultured with 293 cells under the indicated experimental conditions. **** p < 0.0001.

In contrast to CHAF1A and CHAF1B, HIRA mRNA and protein abundance was not significantly changed by primary human B-cell EBV infection, perhaps suggesting that HIRA is well positioned to regulate incoming EBV genomes (Fig. 5F and S6B). We therefore tested whether HIRA also had a role in latency establishment in newly-infected Akata cells. HIRA sgRNA expression increased the percentage of gp350+ cells amongst newly infected GFP+ Akata (Fig. 5G-H), albeit less robustly than CHAF1B sgRNA. Addition of acyclovir strongly reduced the percentage of gp350+ cells, suggesting that lytic replication drove its expression in the context of HIRA depletion. Thus, our results are consistent with a model in which HIRA and CAF1 have non-redundant roles in regulation of EBV latency in Burkitt cells.

The histone H3.3 loaders ATRX and DAXX have roles in telomeres and have been implicated in maintenance of Burkitt B-cell EBV latency. ShRNA targeting of either ATRX or DAXX induces lytic antigen expression (23). Consistent with these RNAi results, we found that CRISPR targeting of either ATRX or DAXX induced BZLF1 and BMRF1 expression on the mRNA and protein levels, but more weakly induced plasma membrane gp350 expression (Fig. S7). Whereas sgRNAs targeting DAXX induced ~2.5 fold increases in EBV copy number, ATRX sgRNAs failed to do so (Fig. S7). Collectively, these data indicate that multiple histone loaders have non-redundant roles in maintenance of EBV latency.

### Loss of CHAF1B reduces the occupancy of H3.1 and H3.3 at EBV lytic genes’ promoters

CAF1 preferentially loads H3.1/H4 histone tetramers onto newly synthesized or damaged host DNA, though whether it is important for H3.1 loading onto latent EBV genomes remains unknown. In addition, little is presently known about whether histone H3.1 versus 3.3 occupancy at key EBV genomic sites in latency. We therefore used chromatin immunoprecipitation (ChIP) for endogenous histone 3.1 and qPCR to investigate effects of CHAF1B depletion on histone 3.1 occupancy at key EBV genomic sites. For cross-comparison, ChIP for histone 3.3 was also performed in parallel on the same samples.

CHAF1B depletion significantly decreased histone 3.1 (H3.1) occupancy at the immediate early *BZLF1* promoter, and at the late gene *BLLF1* (encodes gp350) promoter. Likewise, sgCHAF1B expression decreased H3.1 occupancy at both origins of lytic replication (*oriLyt* L and R), which are EBV genomic enhancers with key roles in lytic gene induction and in lytic DNA replication(52–54) (Fig. 6A). Similar results were obtained in acyclovir-treated cells, suggesting that production of unchromatinized lytic genomes did not falsely lower the ChIP-qPCR result (Fig. S8). This data suggests that latent EBV genomes may be broadly occupied by H3.1-containing nucleosomes, most likely loaded in a DNA replication dependent manner in S-phase (17, 55). Furthermore, CHAF1B depletion reduced H3.3 levels at *BZLF1, BLLF1* and *oriLyt* sites, suggesting that CAF1 also directly or indirectly controls H3.3 loading onto latent EBV genomes. With regards to the latter possibility, RNAseq analysis demonstrated that sgCHAF1B expression diminished the expression of histone and histone-like genes, and ATRX transcript by ~30%, but modestly increased DAXX and HIRA mRNA levels (Table S1). We also note that upon lytic induction by CHAF1B depletion, EBV early gene product BNLF1 is expressed and targets ATRX and DAXX to PML bodies, which may serve to diminish H3.3 loading at these EBV genomic sites (22, 23, 56). Although CHAF1B depletion diminished expression of multiple histone and histone-related genes (Fig. 2F-G and S3A), sgCHAF1B expression did not reduce the steady state H3.1 or H3.3 levels in EBV+ Akata cells, as judged by immunoblot analysis (Fig. 6B).

**Figure 6.**
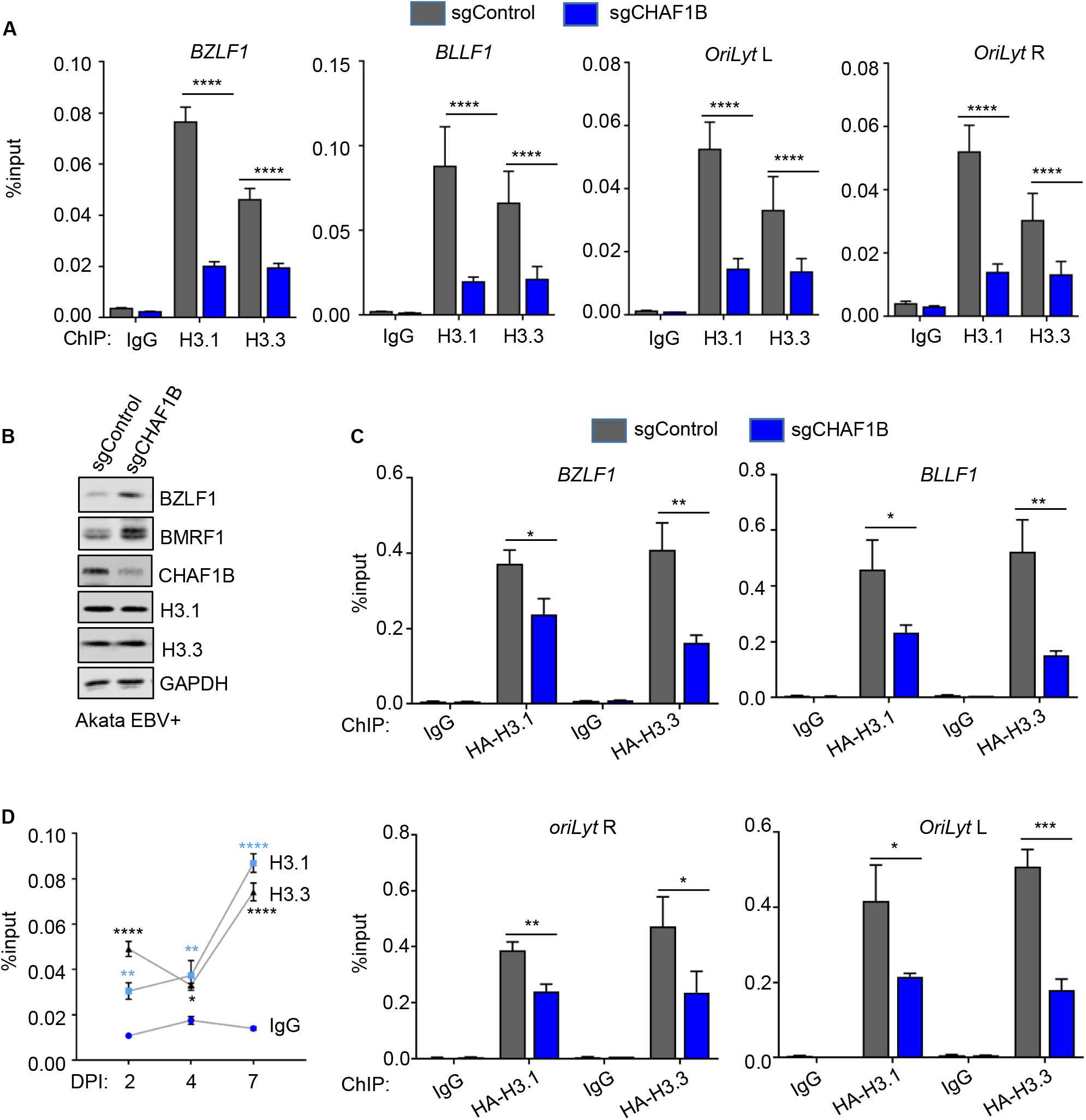
CHAF1B depletion reduces histone H3.1 and H3.3 occupancy at key EBV lytic cycle regulatory elements. (A) ChIP was performed using antibodies against endogenous H3.1 or H3.3 on chromatin from Akata EBV+ cells expressing control or CHAF1B sgRNAs, followed by qPCR with primers specific for the *BZLF1* or *BLLF1* promoters, *oriLyt* R or *oriLyt* L. Mean ± SEM are shown for n=3 biologically independent replicates. p-values were calculated by two-way ANOVA with Sidak’s multiple comparisons test. (B) Immunoblot analysis of WCL from EBV+ Akata, BL cells expressing control or independent CHAF1B. (C) ChIP for HA-epitope tagged H3.1 or H3.3 using anti-HA antibody and chromatin from Akata EBV+ cells stably expressing HA-H3.1 or HA-H3.3 and the indicated sgRNAs. qPCR was then performed with primers specific for the *BZLF1* or *BLLF1* promoters, *oriLyt* R or *oriLyt* L. Mean ± SEM are shown for n=3 biologically independent replicates are shown. p-Values were calculated by two-way ANOVA with Sidak’s multiple comparisons test. (D) ChIP for endogenous H3.1 or H3.3 was performed using antibodies targeting H3.1 or H3.3 on chromatin from human primary B cells infected with B95.8 EBV at 2, 4 and 7 days post infection, followed by qPCR with primers specific for the *BZLF1* promoter. Input DNA for each time point was normalized for intracellular EBV genome copy number. Mean ± SEM are shown for n=3 biologically independent replicates are shown. p-Values were calculated by two-way ANOVA with Sidak’s multiple comparisons test.

To enable additional cross-comparison of CHAF1B perturbation effects on EBV genomic H3.1 and H3.3 occupancy using a single monoclonal antibody, we established EBV+ Akata cells with stable HA-epitope tagged H3.1 or H3.3 expression (49). ChIP was then performed using monoclonal anti-HA antibody in cells expressing sgControl vs sgCHAF1B. Consistent with observations using antibodies against endogenous H3.1 and H.3, CHAF1B depletion similarly reduced HA-3.1 and HA-3.3 signals at *BZLF1, BLLF1*, and *oriLyt* sites (Fig. 6C). These results further suggest that the EBV genome is occupied by H3.1- and H3.3-containing nucleosomes in latency I.

To gain insights into histone H3 isoform loading onto EBV genomes in newly infected primary human B-cells, ChIP-qPCR analyses were performed at 2, 4 and 7 days post EBV-infection. CD19+ B-cells were purified by negative selection and infected with B95.8 EBV at a MOI of 0.2. By Day 2, where infected cells have undergone remodeling but have not yet divided and where most cells should contain only 1-2 EBV genomes (57), H3.1 and H3.3 loading were already significantly increased. This result suggests that both H3 isoforms are loaded onto incoming EBV genomes, potentially by multiple histone loaders. Notably, the EBV tegument protein BNLF1 targets ATRX and DAXX for sequestration in PML bodies at this timepoint (23), suggesting that HIRA and newly induced CAF1 may be responsible. H3.1 and H3.3 levels remained stable at Day 4 post-infection, a timepoint at which cells have entered Burkitt-like hyperproliferation and divide every 8-12 hours (11–13). Interestingly, after the period of Burkitt-like hyper-proliferation that extends roughly from days 3-7 post-infection, H3.1 and H3.3 levels nearly doubled, even when controlling for increases in EBV genome copy number over this interval. This result suggests that each type of histone 3.3 is loaded by host machinery onto newly synthesized episomes, despite short cell cycle times (Fig. 6D).

### CAF1 is important for deposition of repressive H3K9me3 and H3K27me3 heterochromatin marks

CAF1 has important roles in host genome heterochromatin organization (58–60), in part through cross-talk with deposition of repressive histone 3 lysine 9 and 27 trimethyl marks (H3K9me3 and H3K27me3). For instance, in cell fate determination, depletion of CHAF1A reduced H3K27me3 levels at promoters of many genes associated with pluripotency (58). Deposition of H3K9me3 and H3K27me3 marks onto the EBV genome are important for silencing of the lytic and latency III programs (61–66). However, roles its potential roles in regulation of repressive EBV genome repressive H3 marks has not been investigated.

CRISPR knockout was used to test the effects of CHAF1B depletion on H3K9me3 and H3k27me3 marks at four EBV genome lytic cycle sites known to carry these repressive marks in latent B-cell lines. Following expression of control vs CHAF1B sgRNAs in EBV+ Akata cells, ChIP was performed with control IgG or with antibodies against H3K9me3 or H3K27me3. qPCR analysis demonstrated that CHAF1B depletion significantly reduced H3K9me3 occupancy by two to three-fold at the *BZLF1* and *BLLF1* promoters and at both *oriLyt* regions (Fig. 7A). Likewise, sgCHAF1B expression diminished repressive H3K27me3 marks at the sites to a similar extent (Fig. 7B). These results are consistent with a model in which CAF1 histone H3/4 loading onto newly replicated or perhaps DNA damaged EBV episomes is important for the subsequent propagation of repressive H3K9 and H3K27 trimethel heterochromatic marks (Fig. 8).

**Figure 7.**
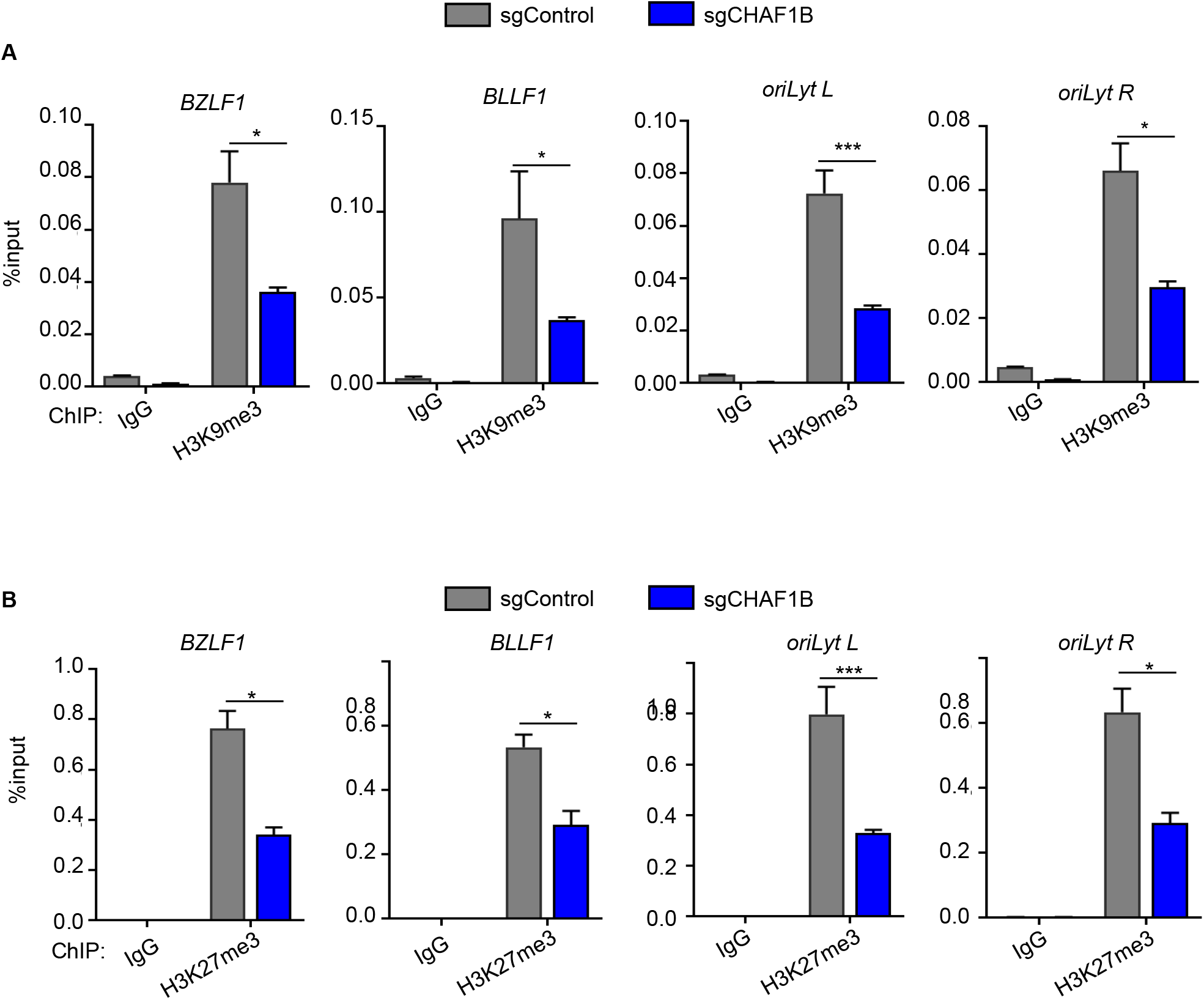
CHAF1b is important for H3K9me3 and H3k27me3 repressive marks at EBV genome lytic cycle regulatory sites. (A) ChIP for H3K9me3 was performed on chromatin from Akata EBV+ cells expressing control or CHAF1B sgRNAs, followed by qPCR with primers specific for the *BZLF1* or *BLLF1* promoters, *oriLyt* R or *oriLyt* L. Mean ± SEM are shown for n=3 biologically independent replicates are shown. p-Values were calculated by two-way ANOVA with Sidak’s multiple comparisons test. (B) ChIP for H3K27me3 was performed on chromatin from Akata EBV+ cells expressing control or CHAF1B sgRNAs, followed by qPCR with primers specific for the *BZLF1* or *BLLF1* promoters, *oriLyt* R or *oriLyt* L. Mean ± SEM are shown for n=3 biologically independent replicates are shown. p-Values were calculated by two-way ANOVA with Sidak’s multiple comparisons test.

**Figure 8.**
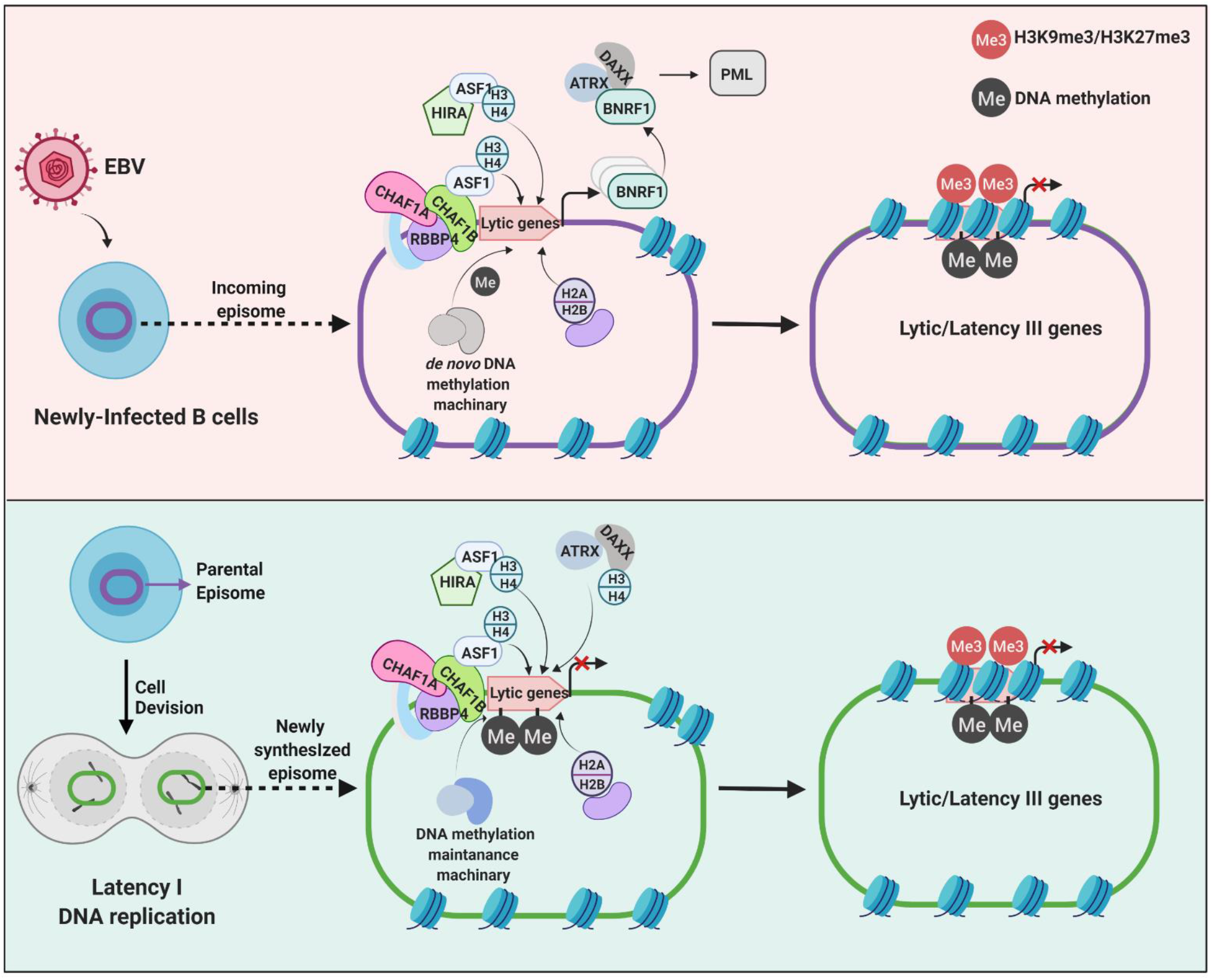
Schematic of histone loader roles in EBV genome regulation. Top, CAF1 and HIRA load histones H3/H4 onto incoming EBV genomes, together with ASF1. H2A/H2B are loaded onto the EBV genome by distinct histone chaperone and assemble into histone octamers. EBV BNRF1 subverts ATRX/DAXX in newly infected cells. DNA methyltransferases and histone H3K9 and H3K27 methyltransferases add repressive marks that suppress lytic cycle and latency III genes. Bottom, CAF1 and HIRA roles in maintenance of latency I. EBV genomes are replicated by host machinery in early S-phase, and newly synthesized genomes must be reprogrammed to latency I. CAF1, HIRA and ATRX/DAXX have non-redundant roles in maintenance of latency I. Cross-talk with DNA methylation machinery is important for propagation of CpG methylation marks that maintain latency I.

## DISCUSSION

EBV coopts host epigenetic pathways to regulate viral genome programs. Incoming EBV genomes are organized into nucleosomes, which must then be maintained or remodeled on newly synthesized, damaged or transcribed regions of EBV genomes. Burkitt lymphoma are amongst the fastest growing human tumor cells, and newly EBV-infected B-cells undergo Burkitt-like hyperproliferation between days 3-7 post-infection in cell culture (11–13). Host machinery must therefore propagate chromatin-encoded epigenetic information with each cell cycle, which begins with histone loading. The results presented here suggest that EBV coopts the CAF1 complex to establish and maintain latency, reminiscent of its use by host pathways that regulate embryonic development and cell fate.

Histone H3 was loaded onto EBV genomes by 48 hours post primary B-cell infection, by which time CAF1 expression was upregulated (35, 36, 67, 68). These observations raise the question of whether CAF1 participates in chromatin assembly on incoming viral genomes. While CRISPR technical limitations currently prevent us from asking this question in resting primary B-cells, our Akata cell system suggested that CAF1 may play a key role in latency establishment. However, in contrast to newly infected primary cells, Akata are rapidly dividing, and our experiments did not differentiate between disruption of latency establishment versus reactivation after the first cell cycle, perhaps related to defects in DNA replication-coupled histone loading. It is plausible that other histone chaperones, in particular HIRA, may play key roles in H3/H4 loading onto incoming EBV genomes prior to mitosis in newly infected cells. CAF1 may then carry out DNA replication dependent roles, beginning with entry of the newly infected cell into S-phase approximately 72 hours post-infection (11–13), and HIRA plays ongoing roles that remain to be defined. Notably, the EBV tegument protein BNLF1 subverts DAXX/ATRX-mediated H3.3 loading on viral chromatin for the first several days post-infection (22, 23) (Fig. 8), but subsequently also become necessary for maintenance of EBV latency, suggesting a complex interplay between multiple histone chaperones.

Histones 3.1 and 3.3 are loaded onto the beta-herpesvirus cytomegalovirus genomes (50), and intriguingly, their deposition did not require transcription or replication of the viral genome. This finding raises the possibility that conserved mechanisms may load histones onto herpesvirus genomes more broadly. Histone 3.1 and 3.3 loading regulate key aspects of herpes simplex gene regulation (49, 69, 70).

The histone chaperone ASF1 transports H3/H4 complexes to the nucleus for deposition onto DNA by CAF1 or HIRA (71). While ASF1a preferentially associates with HIRA and ASF1b with CAF1, they can function redundantly when co-expressed. Depletion of both ASF1A and ASF1B is required to arrest human cell DNA replication (72). Since EBV B-cell infection upregulates both ASF1a and ASF1b transcripts (36), and since ASF1a/b are highly co-expressed in Burkitt cells(73), we speculate that this redundancy precluded either ASF paralogue from scoring in our latency reversal CRISPR screen (24).

Further studies are required to identify how HIRA, ATRX and DAXX maintain latency I, but permit EBNA1 and EBV non-coding RNA expression. It also remains possible that they indirectly control EBV genome expression through effects host transcription factor expression, which then secondarily regulate the EBV genome. Further underscoring the intricate relationship between EBV and histone biology, EBNA3C downregulates the histone H2A variant H2AX shortly after primary B-cell infection at the mRNA and protein levels (74).

MYC suppresses EBV reactivation by preventing DNA looping of *oriLYT* and terminal repeat regions to the *BZLF1* promoter (24). The results presented here raise the question of whether histone loading by CAF1 or HIRA may act together with MYC to prevent long-range EBV genomic DNA interactions that promote lytic reactivation, perhaps at the level of CTCF, cohesion or other DNA looping factors. Perturbation of histone loading may alternatively be sufficient to de-repress *BZLF1*. Indeed, micrococcal nuclease digestion experiments demonstrated that immediate early *BZLF1* and *BRLF1* promoters are nucleosomal (75). Yet, open chromatin at *BZLF1* and *BRLF1* is not sufficient for lytic reactivation (65). Furthermore, CHAF1B de-repression strongly de-repressed latency III gene expression, suggesting a broader role in silencing of EBV antigens.

We recently identified that the facilitated chromatin transcription (FACT) histone loading complex is critical for EBV Burkitt latency (24). FACT remodels histones at sites of active transcription to enable RNA polymerase processivity (76, 77). Further underscoring diverse histone chaperone roles in maintenance of EBV latency, FACT was found to regulate EBV latency through effects on *MYC* expression, consistent with its role in driving glioblastoma oncogenic *N-MYC* expression (78). However, we observed only modest reduction in *MYC* mRNA expression upon CHAF1B knockdown (Table 1), suggesting an alternative mechanism for its EBV latency maintenance role.

A longstanding question has remained how lytic EBV genomes destined for packaging into viral particles evade histone loading, since histones are not detectable in purified EBV viral particles (79). EBV lytic replication is initiated in early S-phase, taking place in nuclear factories that are devoid of histones or host DNA. CAF1 is recruited to host DNA replication forks through association with the DNA clamp PCNA. While PCNA can be detected in EBV amplification factories, it does not localize to sites of viral DNA synthesis (16). Abundances of CHAF1A, CHAF1B, ASF1a and ASF1b decline in lytic replication in Burkitt/epithelial cell somatic hybrid D98/HR1 cells, whereas HIRA and DAXX levels were stable.

DNA methylation is important for suppression of latency III, raising the question of how CHAF1B depletion de-repressed latency III genes (Figure S3A). While we note that latency III transcripts are induced in Burkitt cell EBV lytic reactivation (27), we speculate that CAF1 depletion may perturb maintenance of EBV genomic DNA methylation through effects on crosstalk between histone and DNA methylation pathways. ChIP-seq approaches demonstrated H3K9me3 and H3K27me3 repressive marks at key lytic and latency gene sites (61, 80, 81). Furthermore, we recently found domains of the enzyme UHRF1 that read H3K9me2/me3, the H3 N-terminus and hemi-methylated DNA are essential for EBV latency I (73). We therefore speculate that CHAF1B depletion may perturb UHRF1 recruitment and thereby disrupt DNA methylation.

CHAF1B depletion induced a strong interferon induced signature, which we and other have not observed in Burkitt cell lytic reactivation triggered by immunoglobulin cross-linking or by conditional BZLF1 alleles. Therefore, we hypothesize that DNA sensing pathways may be activated by CHAF1B depletion, for example in response to exposure of viral or host non-chromatinized DNA. Alternatively, latency III triggers interferon induced genes, and a de-repressed EBV transcript may be responsible for this phenotype (35, 36, 82). It is also worth noting that CHAF1B depletion resulted in downregulation of numerous histone and histone-like genes, which we speculate may result from a negative feedback loop that responds to loss of this important histone chaperone complex.

EBV exclusively establishes infection in normal, differentiated epithelial cells (83–86). Epithelial cell replication plays important roles in EBV shedding into saliva (87), and uncontrolled lytic EBV replication can cause oral hairy leukoplakia in heavily immunosuppressed people. It will be of significant interest to determine how CAF1, HIRA, ATRX and DAXX roles may be distinct in epithelial cells to support escape from EBV latency.

Current Burkitt lymphoma therapies cause major side effects and increase the risk of secondary malignancies. eBL management is further complicated by the risk of giving high-intensity chemotherapy in resource-limited settings. Consequently, there is significant interest in developing safer therapeutic regimen, including EBV lytic reactivation strategies (88). Reversal of EBV Burkitt latency could selectively sensitize tumor cells to T-cell responses and to the antiviral drug ganciclovir (89).

## Supplemental Figure Legends

**Figure S1.**
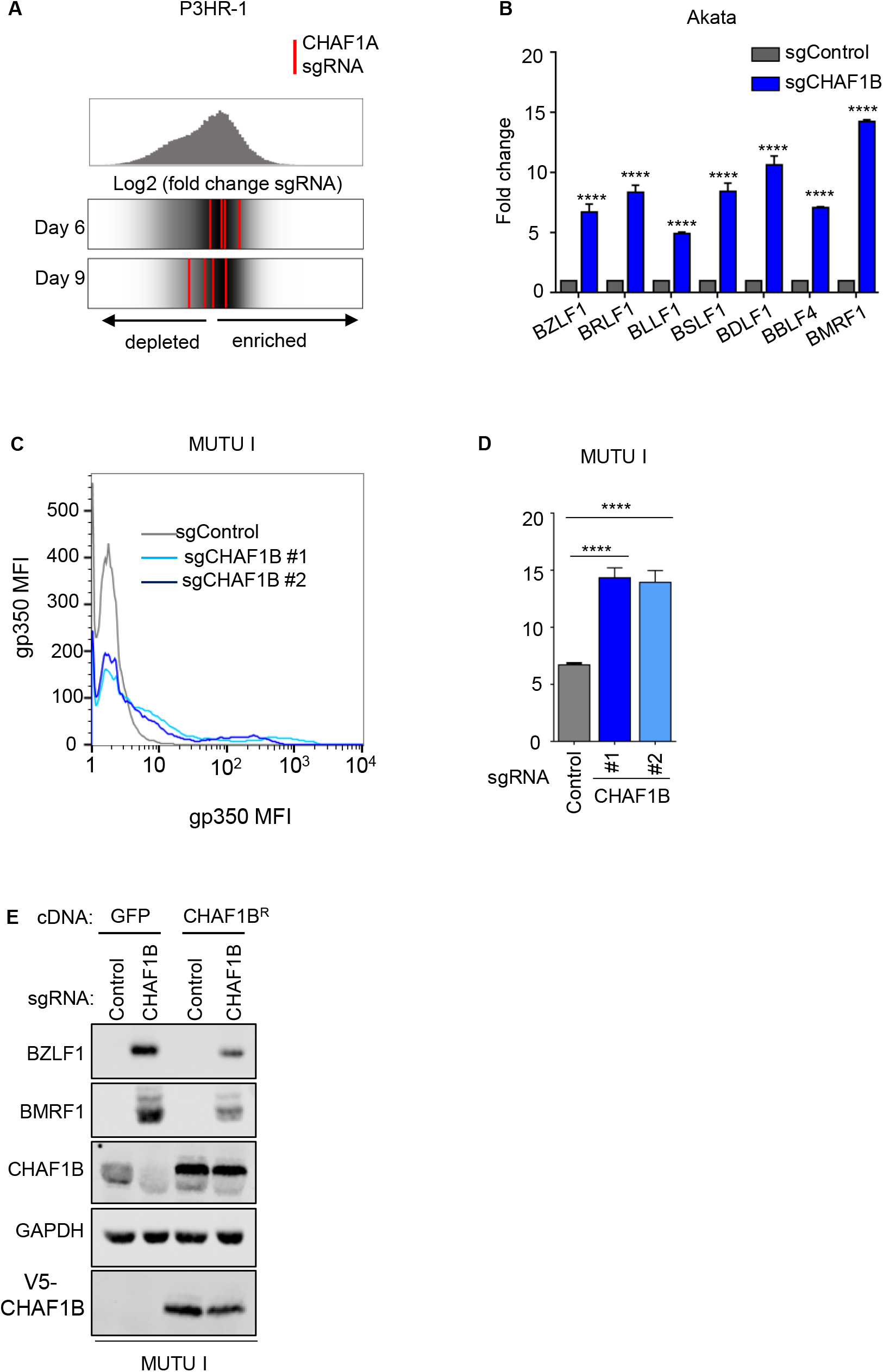
CHAF1B depletion triggers EBV lytic antigens expression in Burkitt cells. (A) Top: Distribution of Log2 fold-change (LFC) values of sgRNAs in gp350+ sorted versus input library cells for all Avana library guides at screen Day 6. Bottom: LFC for the four *CHAF1A* targeting sgRNAs (red lines), overlaid on gray gradient depicting the overall sgRNA distribution, at CRISPR screen Days 6 versus 9. Average values from two screen biological replicates are shown. (B) qRT-PCR analysis of selected viral immediate early, early, and late genes in Akata EBV+ cells expressing control or independent CHAF1B sgRNA. Mean + SD values from N=3 replicates, **** p < 0.0001 (C) FACS analysis of PM gp350 expression in MUTU I cells expressing control or independent CHAF1B sgRNAs. (D) Mean + SD PM gp350 MFI values from n=3 replicates of Akata with indicated sgRNAs, as in (C). **** p < 0.0001. (E) Immunoblot analysis of WCL from MUTU I cells stably expressing GFP or V5 epitope tagged-CHAF1B cDNAs and control or CHAF1B sgRNA, as indicated. Blot is representative of n=3 replicates.

**Figure S2.**
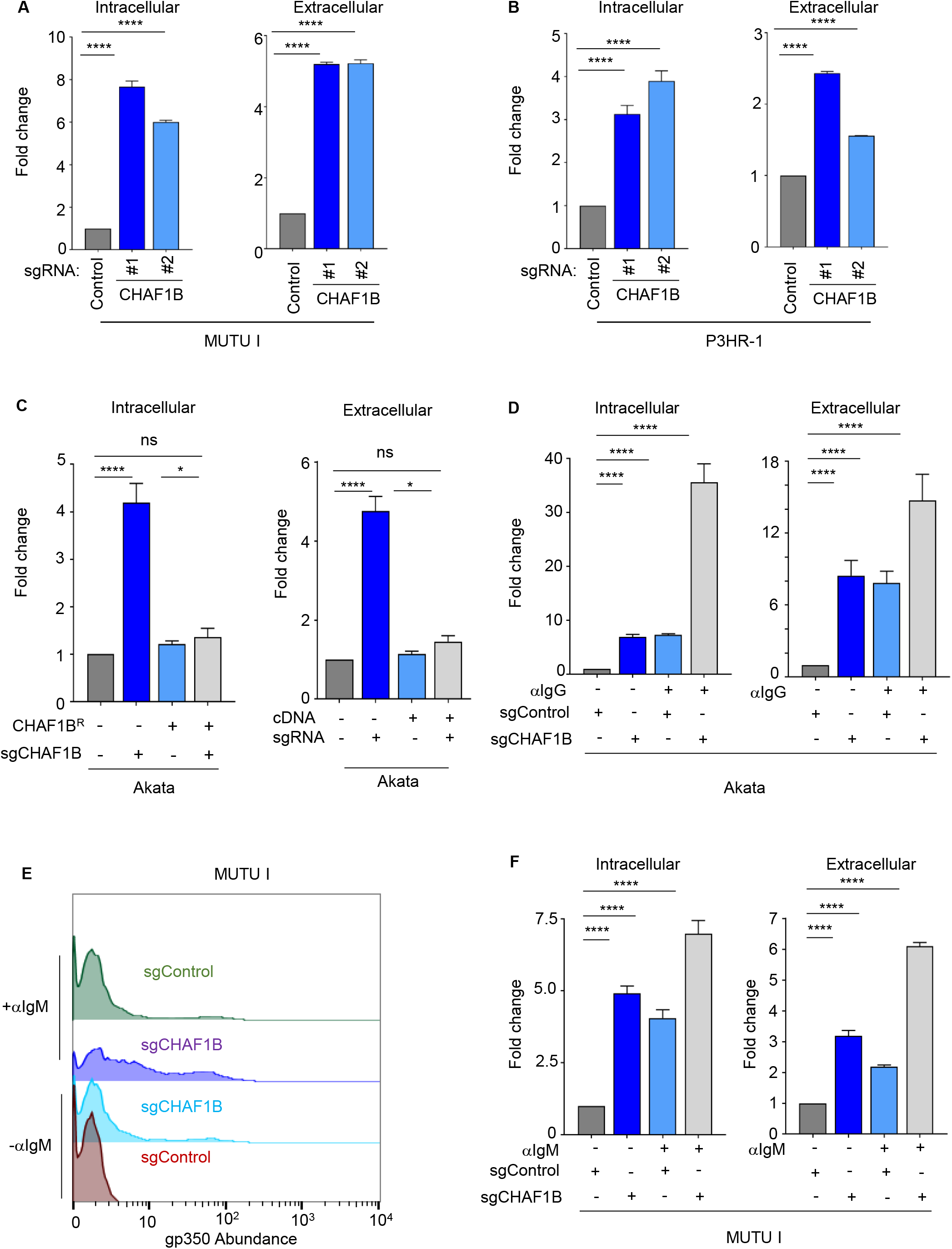
Depletion of CHAF1B induces lytic reactivation in multiple EBV+ Burkitt tumor cell lines. (A, B) qPCR analysis of EBV intracellular genome copy number from MUTU I (A) or P3HR-1 (B) cells expressing control or CHAF1B sgRNAs. Total genomic DNA was extracted at Day 6 post-lentivirus transduction. Mean ±SD values from n=3 replicates are shown. ****p<0.0001. (C) qPCR analysis of EBV intracellular or extracellular genome copy number from Akata EBV+ cells expressing GFP of V5-CHAF1B^R^ cDNAs and the indicated sgRNAs. Cells expressed GFP where not indicated to express CHAF1B^R^, and cells expressed sgControl where not indicated to express sgCHAF1B. Total genomic DNA was extracted at Day 6 post lentivirus transduction. Mean ± SD values from n=3 replicates are shown. ****p<0.0001, *<0.05, ns=non-significant. (D) qPCR analysis of EBV intracellular or extracellular genome copy number from Akata cells expressing control or CHAF1B sgRNAs comparing alone or in combination with 10μg/ml of αIgG for 48 hours. Mean ±SD values from n=3 replicates are shown. ****p<0.0001. (E) FACS analysis of PM gp350 expression in MUTU I cells expressing control or independent CHAF1B sgRNAs, and mock-induced or induced with αIgM 10μg/ml. (F) qPCR analysis of EBV intracellular or extracellular genome copy number from MUTU I expressing control or CHAF1B sgRNAs alone or in combination with αIgM induction. KO cells were mock-treated or treated with 10μg/ml of αIgM for 48 hours. Mean ±SD values from n=3 replicates are shown. ****p<0.0001.

**Figure S3.**
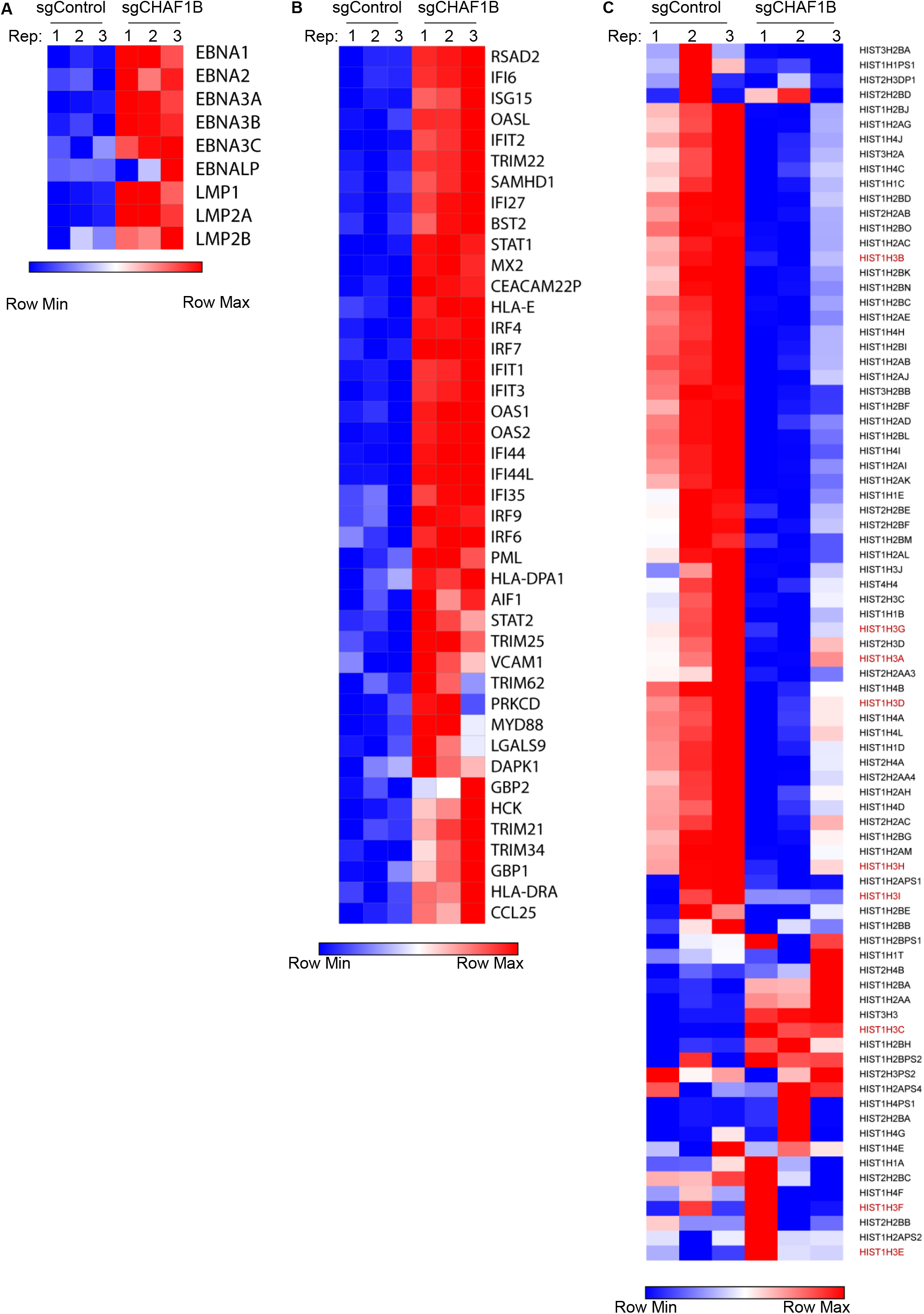
RNAseq heatmap analyses of CHAF1 depletion effects on Akata EBV latency III, interferon stimulated gene and histone gene expression. (A) Heatmap representation of EBV latency III gene abundance in Akata EBV+ cells expressing control or CHAF1B sgRNAs. Shown are data from n=3 biologically independent replicates. (B) Heatmap representation of interferon stimulated genes whose expression was significantly upregulated in sgCHAF1b versus sgControl expressing Akata EBV+ cells. Shown are data from n=3 replicates. (C) Heatmap representation of mRNAs encoding histones whose expression was significantly different in Akata EBV+ cells expressing sgCHAF1B vs sgControl. Shown are data from n=3 replicates. Histone H3 genes are labeled in red.

**Figure S4.**
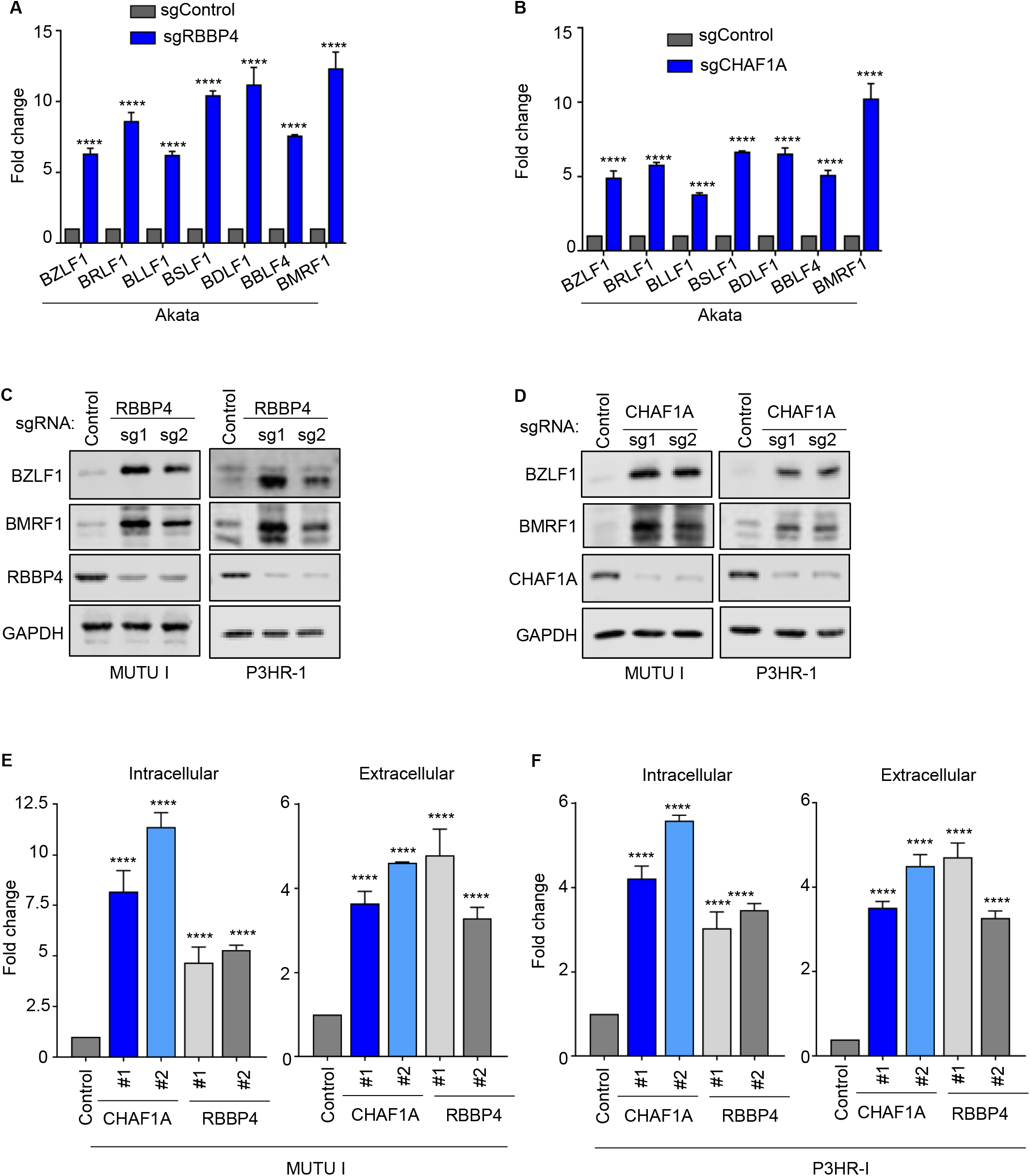
CHAF1A or RBBP4 depletion trigger Burkitt cell EBV lytic reactivation. (A and B) qRT-PCR analysis of selected viral immediate early, early, and late genes from Akata EBV+ cells expressing control or independent RBBP4 (A) or CHAF1A (B) sgRNAs. Mean ±SD values from n=3 replicates are shown. ****p<0.0001 (C and D) Immunoblot analysis of WCL from MUTU I or P3HR-1 cells expressing control or independent RBBP4 (C) or CHAF1A (D) sgRNAs. Blots are representative of n=3 replicates. (E) qPCR analysis of EBV intracellular or extracellular genome copy number from MUTU I cells expressing control, CHAF1A or RBBP4 sgRNAs. Mean ±SD values from n=3 replicates are shown. ****p<0.0001, ***p<0.001. (F) qPCR analysis of EBV intracellular or extracellular genome copy number from P3HR-1 cells expressing control, CHAF1A or RBBP4 sgRNAs. Mean ±SD values from n=3 replicates are shown. ****p<0.0001, ***p<0.001.

**Figure S5.**
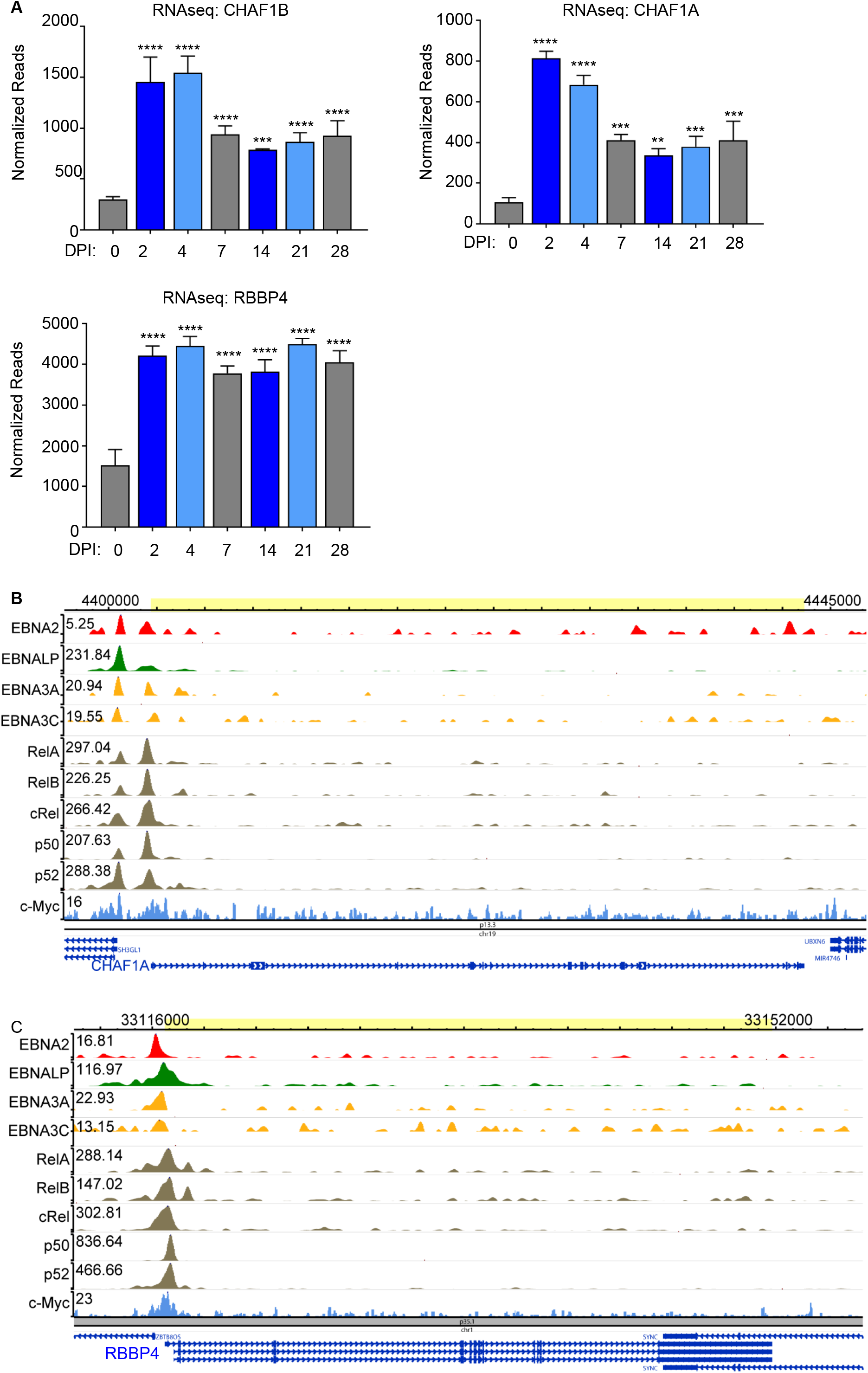
EBV induces CAF1 in newly infected primary human B-cells. (A) Normalized CHAF1B, CHAF1A, or RBBP4 mRNA levels from primary human peripheral blood B-cells at the indicated day post infection (DPI) by the EBV B95.8 strain (36). Shown are the mean ± SEM values from n=3 of biologically independent RNAseq datasets. ****p<0.0001, ***p<0.001, **p<0.01. (B and C) LCL ChIP-seq signals of EBNA2, EBNALP, EBNA3A, 3C, RelA, RelB, cRel, p50, p52 and MYC at the *CHAF1A* (B) or *RBBP4* (C) loci. Track heights are indicated in the upper left, and genomic positions indicated at top of each panel.

**Figure S6.**
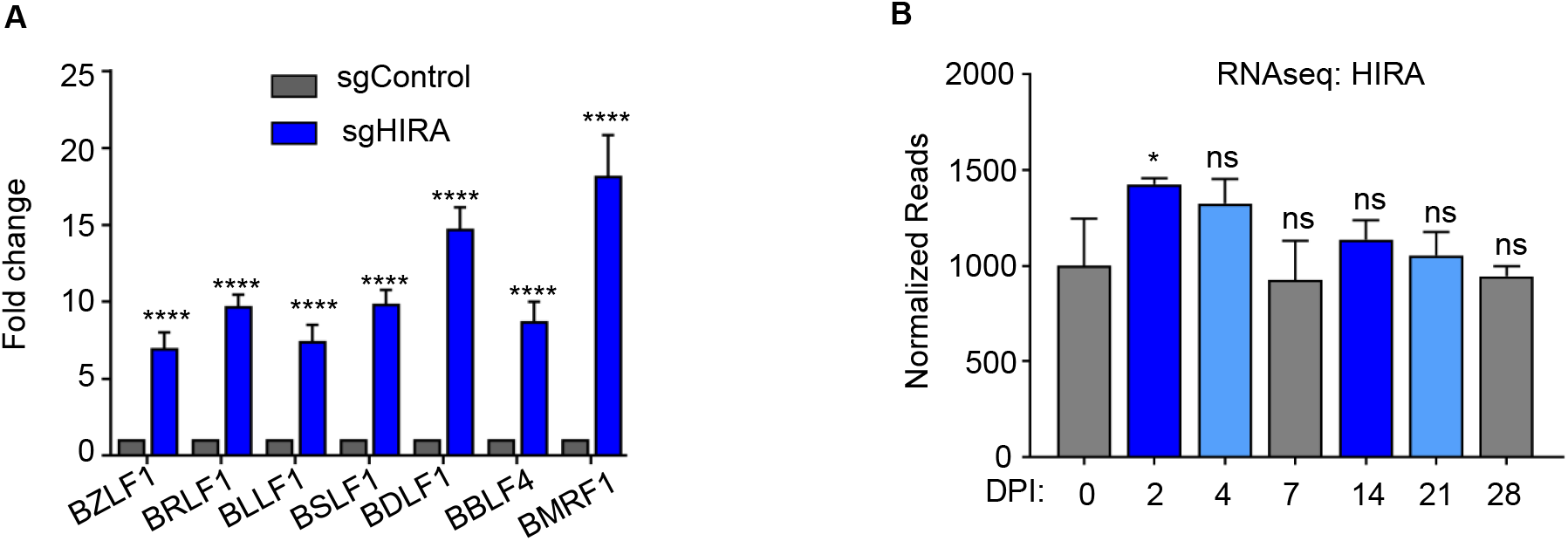
HIRA depletion triggers Burkitt cell EBV lytic reactivation. (A) qRT-PCR analysis of selected viral immediate early, early, and late genes from Akata EBV+ cells expressing control or HIRA sgRNAs. Mean + SD shown from n=3 replicates, **** p<0.0001. (B) Normalized HIRA mRNA levels in primary human peripheral blood B-cells at the indicated day post infection (DPI) by EBV B95.8 (36). Shown are the mean + SD values from n=3 of biologically independent RNAseq replicates. *, p<0.05. ns, non-significant.

**Figure S7.**
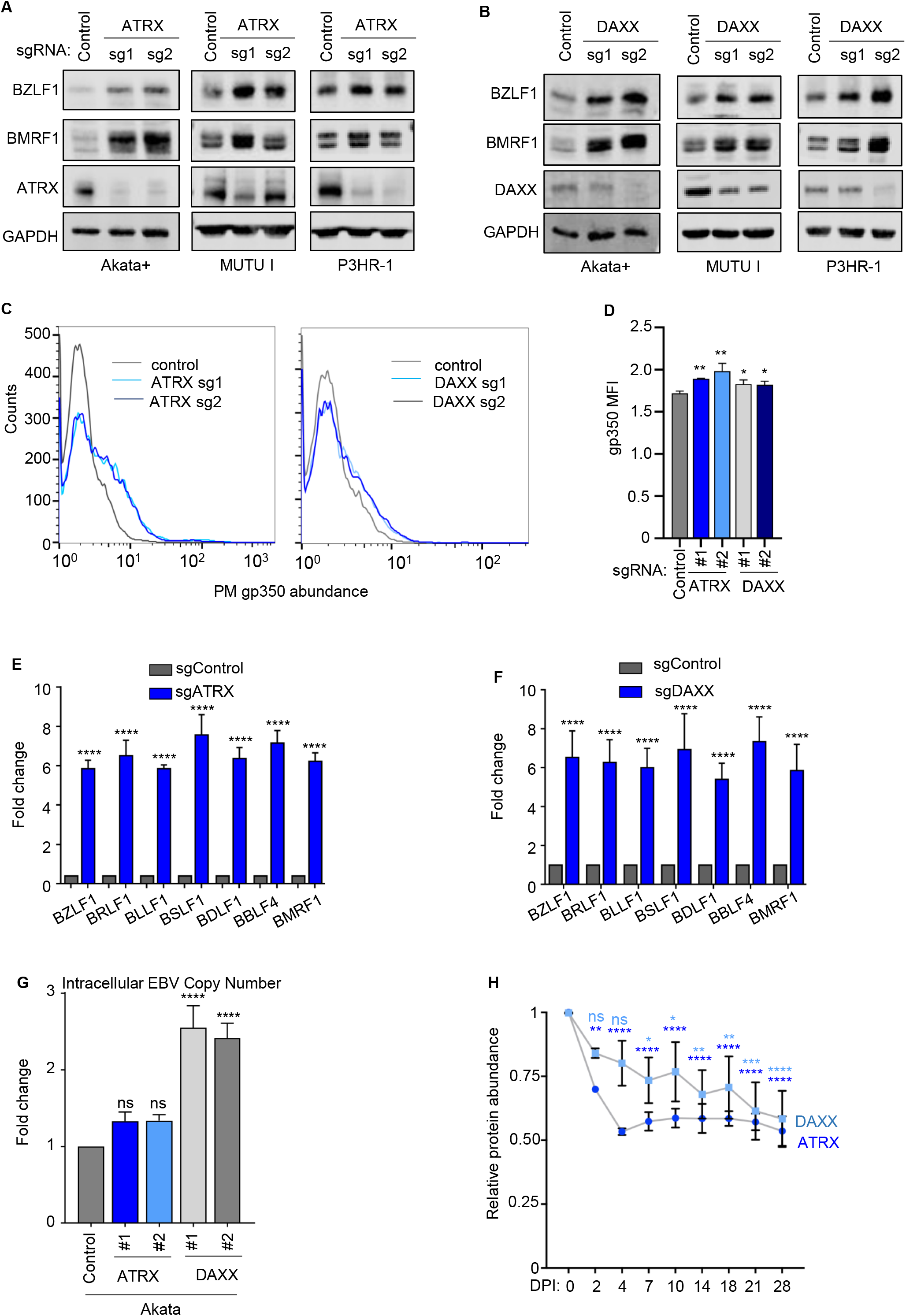
DAXX and ATRX depletion triggers Burkitt EBV lytic reactivation. (A) Immunoblot analysis of WCL from EBV+ Akata, MUTU I and P3HR-1 cells expressing control or independent ATRX sgRNAs. (B) Immunoblot analysis of cell extracts from EBV+ Akata, MUTU I and P3HR-1 cells expressing control, or DAXX sgRNAs. (C) FACS analysis of PM gp350 expression in Akata EBV positive cells expressing control, ATRX, or DAXX sgRNAs. (D) Mean ± SD PM gp350 MFI values from n=3 replicates of Akata EBV positive cells expressing control, ATRX, or DAXX sgRNAs, as in (C). **** p < 0.0001. (E and F) qRT-PCR analysis of selected viral immediate early, early, and late genes in Akata EBV+ cells expressing control or independent ATRX(A) or DAXX (E) sgRNAs. Mean + SD from n=3 replicates, **** p<0.0001. (G) qPCR analysis of EBV intracellular genome copy number from Akata EBV+ cells expressing control, ATRX or DAXX sgRNAs. Mean +SD values from n=3 replicates are shown. ****p<0.0001, ns=non-significant. (H) ATRX or DAXX relative protein abundances detected by tandem-mass-tag-based proteomic analysis of primary human B-cells at rest and at nine time points after EBV B95.8 infection at a multiplicity of infection of 0.1. Data represent the average +/- SEM for n=3 independent replicates(35). For each protein, the maximum level detected across the time course was set to a value of one. Blots in A, B are representative of n=3 replicates.

**Figure S8.**
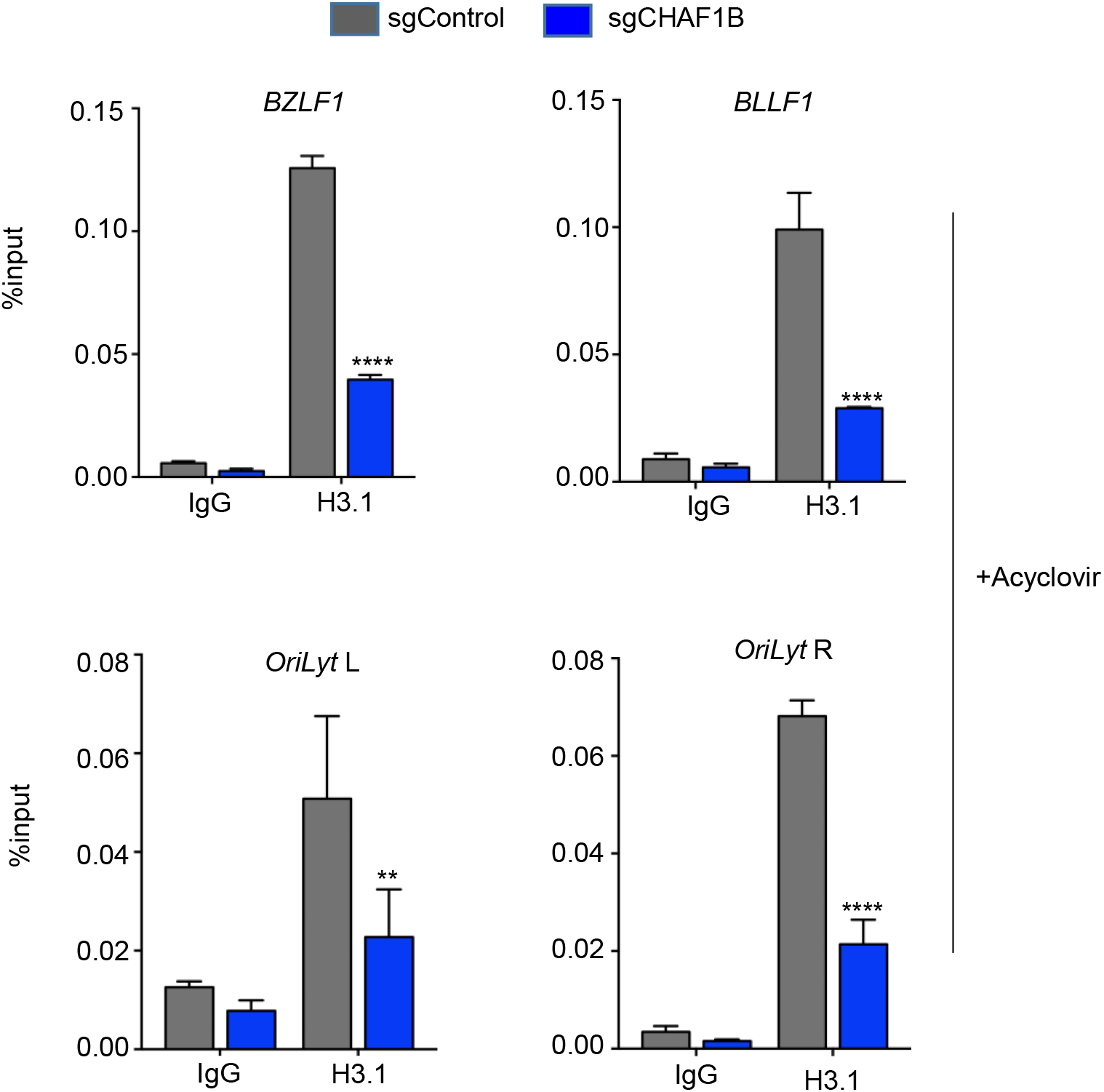
CHAF1B depletion reduces H3.1 loading at multiple EBV genomic lytic cycle regulatory elements in presence of acyclovir. (A) ChIP for H3.1 or H3.3 was performed using antibodies targeting endogenous H3.1 or H3.3 on chromatin from Akata EBV+ cells expressing control or CHAF1B sgRNAs, treated with 100μg/ml acyclovir. qPCR with primers specific for *BZLF1* or *BLLF1* promoters, *oriLyt* R or *oriLyt* L. Mean ± SEM are shown for n=3 biologically independent replicates are shown, ****p<0.0001, **p<0.01. p-Values were calculated by two-way ANOVA with Sidak’s multiple comparisons test.

## Supplementary Tables

**Table S1** Differentially expressed genes in EBV+ Akata cells expressing control or CHAF1B sgRNAs and Enrichr analysis of selected genes (p<0.05, LFC>1 or <−1)

**Table S2** List of antibodies, reagents, kits, and oligoes used in this study.

## MATERIALS AND METHODS

### Cell lines and culture

Throughout the manuscript, all B-cell lines used stably expressed *S. pyogenes* Cas9. The EBV+ Burkitt lymphoma cell lines P3HR-1, Akata, and MUTU I were used in the study. EBV-Akata cells are a derivative cell line of the original EBV+ Akata tumor cell line that spontaneously lost EBV in culture. The EBV+ Burkitt lymphoma cell lines Akata EBV+, MUTU I, P3HR-1 and EBV-Akata were maintained in RMPI 1640 (Gibco, Life Technologies) supplemented with 10% fetal bovine serum (Gibco). 293T were grown in Dulbecco’s Modified Eagle’s Medium (DMEM) with 10% fetal bovine serum (Gibco). Cell lines with stable expression of *Streptococcus pyogenes* Cas9 gene were generated by lentiviral transduction, followed by blasticidin selection at 5 μg/ml, as reported (90). For selection of transduced cells, puromycin was added at the concentration of 3 μg/ml. Hygromycin was used at 200 μg/ml for the initial 4 days, and 100 μg/ml thereafter. Acyclovir was used at the concentration of 100 μg/ml in vitro. EBV producer HEK-293 cells stably transformed by BART-repaired B95-8 based EBV BAC system encoding GFP (45) were cultured in RPMI 1640 (Gibco, Life Technologies) supplemented with 10% fetal bovine serum (Gibco) and 1% penicillin, 50 μg/ml hygromycin. All cells used in this study were cultured in a humidified incubator at 37°C with 5% CO2 and routinely tested and certified as mycoplasma-free using the MycoAlert kit (Lonza). STR analysis (Idexx) was done to verify identify of MUTU I cells.

### Immunoblot analysis

Immunoblot analysis was performed as previously described (91). In brief, WCL were separated by SDS-PAGE electrophoresis, transferred onto nitrocellulose membranes, blocked with 5% milk in TBST buffer and then probed with primary antibodies at 4 °C overnight on a rocking platform, washed four times and then incubated with secondary antibody (Cell Signaling Technology, cat#7074 and cat#7076) for 1 h at room temperature. Blots were then developed by incubation with ECL chemiluminescence for 1 min (Millipore, cat#WBLUF0500) and images were captured by Licor Fc platform. All antibodies used in this study were listed in supplementary Table S2.

### Flow cytometry analysis

For live cells staining, 1× 10^6^ of cells were washed twice with FACS buffer (PBS, 1mM EDTA, and 0.5% BSA), followed by primary antibodies incubation for 30 min on ice. Labeled cells were then washed three times with FACS buffer. Data were recorded with a BD FACS Calibur and analyzed with Flowjo X software (Flowjo).

### Quantification of EBV genome copy number

To measure EBV genome copy number, intracellular viral DNA and virion-associated DNA present in cell culture supernatant were quantitated by qPCR analysis. For intracellular viral DNA extraction, total DNA from 2×10^6^ of Burkitt cells was extracted by the Blood & Cell culture DNA mini kit (Qiagen #13362). For extracellular viral DNA extraction, 500 μl of culture supernatant was collected from the same experiment as intracellular DNA measurement, and was treated with 20 μl RQ1 DNase (Promega) for 1 h at 37°C to degrade non-encapsidated EBV genomes. 30 μl proteinase K (20 mg/ml, New England Biolabs, #P8107S) and 100 μl 10% (wt/vol) SDS (Invitrogen, #155553-035) were then were added to the reaction mixtures, which were incubated for 1 h at 65°C. DNA was purified by phenol-chloroform extraction followed by isopropanol-sodium acetate precipitation and then resuspended in 50 μl nuclease-free water (Thermo Fisher, #10977-023). Extracted DNA was further diluted to 10 ng/ul and subjected to qPCR targeting of the EBV *BALF5* gene. Standard curves were made by serial dilution of a pHAGE-BALF5 miniprep DNA at 25 ng/uL. Viral DNA copy number was calculated by inputting sample Ct values into the regression equation dictated by the standard curve.

### cDNA rescue assay

V5-tagged CHAF1B cDNA with G360A PAM site mutation was synthesized by Genescript (Piscataway, NJ), as described in the following table. CHAF1B sg1 sequence is shown. PAM sequences are underlined. Mutation site is indicated in red. Rescue cDNA was synthesized by GenScript (Piscataway, NJ) and cloned into pLX-TRC313 vector. Cas9 expressing B cells with stable C-terminal V5 epitope-tagged CHAF1B cDNA expression was established by lentiviral transduction and hygromycin selection.

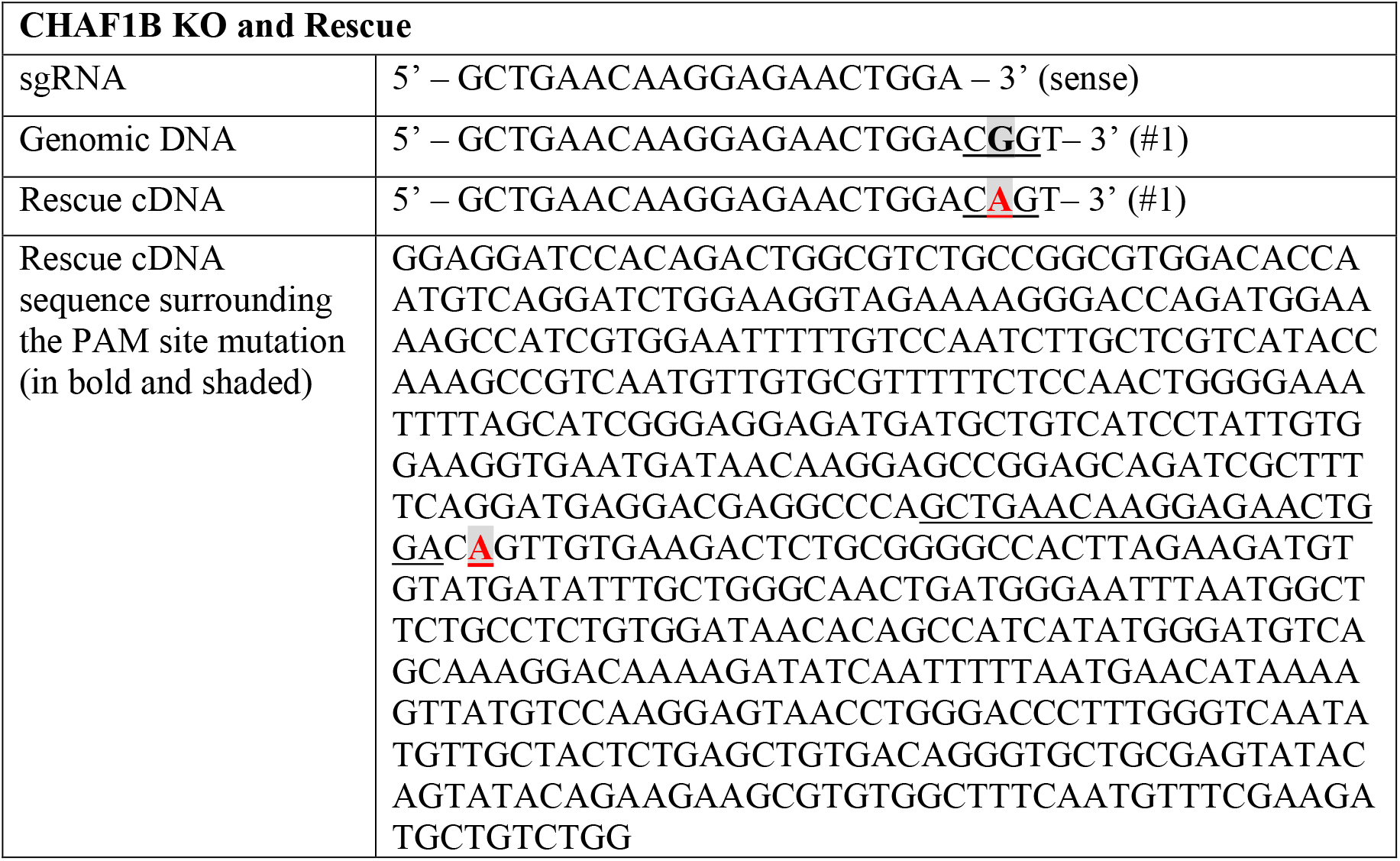

### Chromatin Immunoprecipitation (ChIP) qPCR

Cells were crosslinked with formaldehyde 0.4% for 10 min at room temperature and the reaction was stopped by adding glycine (2.5M) to final concentration 0.2M for 10 minutes at room temperature. The cells were washed three times with PBS and then lysed by 1% SDS lysis buffer (50mM Tris pH8.1, 10mM EDTA, 1% SDS and protease inhibitor) for 20min on ice. Lysate was sonicated 25 min (30 sec on / 30 sec off) in a Diagenode water bath-sonicator and centrifuged at 13000 rpm for 10 min. The supernatant was diluted 10 times in ChIP Dilution Buffer (SDS 0.01%, Triton X-100 1.1%, 1.2 mM EDTA pH 8, 16.7 mM Tris-HCl pH 8.1, 167 mM NaCl and protease inhibitor) and pre-cleared for 1 hour, rotating at 4°C with blocking beads. Soluble chromatin was diluted and incubated with 4μg anti-HA polyclonal antibody (Abcam, #ab9110), anti-H3.1/H3.2 polyclonal antibody (Millipore, #ABE154) or anti-H3.3 polyclonal antibody (Millipore, #09-838). Specific immunocomplexes were precipitated with protein A beads (Thermo fisher, #101041). The beads were washed, for 5 minutes, once in Low Salt Buffer (SDS 0.1%, Triton X-100 1%, 2 mM EDTA pH 8.1, 20 mM Tris-HCl pH 8.1 and 150 mM NaCl), twice in High Salt Buffer (SDS 0.1%, Triton X-100 1%, 2 mM EDTA pH 8, 20 mM Tris-HCl pH 8.1 and 500 mM NaCl), once in LiCl Buffer (0.25 M LiCl, NP-40 1%, Na Deoxycholate 1%, 1 mM EDTA pH 8.1 and 10 mM Tris-HCl pH 8.1) and twice in TE buffer. After reverse cross-linking, DNA was purified by using QIAquick PCR purification kit (Qiagen, #28106). qPCR quantified the DNA from ChIP assay and normalized it to the percent of input DNA. Primers for qPCR are listed in Supplementary Table S2.

### RT-PCR analysis

Total RNA was harvested from cells using RNeasy Mini Kit (Qiagen, #27106). Genomic DNA was removed by using the RNase-Free DNase Set (Qiagen, #79254). RNA was reversed transcribed by iScriptTM Reverse Transcription Supermix (Bio-Rad, #1708841). qRT-PCR was performed using Power SYBR Green PCR Mix (Applied Biosystems, #4367659) on a CFX96 Touch™ Real-Time PCR Detection System (Bio-Rad), and data were normalized to internal control GAPDH. Relative expression was calculated using 2-ΔΔCt method. All samples were run in technical triplicates and at least three independent experiments were performed. The primer sequences were listed in Supplementary Table S2.

### Primary Human B Cells Purification

Discarded, de-identified leukocyte fractions left over from platelet donations were obtained from the Brigham and Women’s Hospital blood bank. Peripheral blood cells were collected from platelet donors, following institutional guidelines. Since these were de-identified samples, the gender was unknown. Our studies on primary human blood cells were approved by the Brigham & Women’s Hospital Institutional Review Board. Primary human B cells were isolated by negative selection using RosetteSep Human B Cell Enrichment and EasySep Human B cell enrichment kits (Stem Cell Technologies, #15064 and #19054), according to the manufacturers’ protocols. B cell purity was confirmed by plasma membrane CD19 positivity through FACS. Cells were then cultured with RPMI 1640 with 10% FBS.

### EBV Infection of Primary B-Cells

EBV B95-8 virus was produced from B95-8 cells with conditional ZTA expression. 4HT was used at a concentration of 1 μM to induce EBV lytic replication, removed 24 hours later, and cells were resuspended in 4HT-free RPMI/10% FBS for 96 hours. Virus-containing supernatants were collected and subject to filtration through a 0.45 μm filter to remove producer cells. Titer was determined experimentally by transformation assay as described previously (35). For analysis of transforming EBV production in Burkitt knockout experiments, culture supernatants from Akata EBV+ cells expressing control, CHAF1B or HIRA sgRNAs were harvested. Supernatants were passed through a 0.80 μm filter to remove any producer cells and were then mixed with 1 million purified CD19+ primary human B cells in 12 well plates. For determining histone H3.1 or H3.3 occupancy in newly infected primary cells, 6×10^7 purified human B cells were infected with B95.8 at a MOI of 0.2. Ten million cells were harvested at 2, 4, and 7 DPI. Viral episome number at each time point was quantitated by *BALF5* qPCR. The recombinant vector pHAGE-BALF5 was used to establish the standard curve for absolute quantification of EBV episome number. The H3 ChIP qPCR signals were normalized using EBV episome numbers at each time point, in order to control for changes in EBV copy number in B-cells between DPI 2-7.

### Co-cultivation of Akata EBV negative cells EBV HEK-293 producer cells

EBV producer HEK-293 cells stably transformed by BART-repaired B95-8 based GFP-EBV BAC system (45). EBV producer cells were seeded at a density of 0.3×10^6/ml in Corning^®^ BioCoat™ Collagen I 6 Well Plate (cat#356400). After 24 hours, HEK-293 producer cells were co-transfected with 500μg of pCDNA-BALF4 and 500ug of pCDNA-BZLF1 per well, as described previously(92).

After incubating for additional 24h, 0.5×10^6/ml of control or CHAF1B KO of Akata EBV-cells resuspended in fresh media were added onto the HEK-293 cells gently. Co-cultured cells were then mock-treated or treated with 100μg/ml Acyclovir (Cat#114798, Millipore). After additional 48 hours, Akata cells were resuspended carefully, without disturbing the 293 monolayer, transferred into a new 6 well plate and further settled for another 24 hours for the removal of potentially contaminating 293 cells. Akata cells were then subjected to the gp350 PM FACS. FSC and SSC parameters were used to exclude any potentially contaminating 293 producer cells.

### RNA sequencing (RNAseq) analysis

Total RNAs were isolated with RNeasy Mini kit using the manufacturer’s protocol. An in-column DNA digestion step was included to remove residual genomic DNA contamination. To construct indexed libraries, 1 μg of total RNA was used for polyA mRNA-selection using NEBNext Poly(A) mRNA Magnetic Isolation Module (New England Biolabs), followed by library construction via NEBNext Ultra RNA Library Prep Kit for Illumina (New England Biolabs). Each experimental treatment was performed in triplicate. Libraries were multi-indexed, pooled and sequenced on an Illumina NextSeq 500 sequencer using single-end 75 bp reads (Illunima).

For RNA-seq data analysis, paired-end reads were mapped to human (GENCODE v28) and the Akata EBV genome. Transcripts were quantified using Salmon v0.8.2 (93) under quasi-mapping and GC bias correction mode. Read count table of human and EBV genes was then normalized across compared cell lines/conditions and differentially expressed genes were evaluated using DESeq2 v1.18.1 (94) under default settings.

Volcano plots were built based on the log2 (foldchange) and −log10 (p-Value) with Graphpad Prism7. Heatmaps were generated by feeding the Z-score values of selected EBV genes from DESeq2 into Morpheus(https://software.broadinstitute.org/morpheus/). Enrichr was employed to perform gene list-based gene set enrichment analysis on selected gene subset(95). Consistent enriched gene sets in Top 5 terms ranked by Enrichr adjusted p-value were visualized Graphpad Prism 7.

### Statistical analysis

Data are presented as mean ± standard errors of the mean. Data were analyzed using analysis of variance (ANOVA) with Sidak’s multiple comparisons test or two-tailed paired Student t test with Prism7 software. For all statistical tests, a cutoff of p < 0.05 was used to indicate significance.

## ACKNOWLEDGEMENTS

This work was supported by NIH RO1s AI137337 and CA228700 (BEG), Starr Cancer Foundation Project # I11-0043, Burroughs Wellcome Career Award in Medical Sciences and American Cancer Society Research Scholar Award (BEG). We thank Joe Cabral and David Knipe for HA-H3.1 and 3.3 expression vectors and for technical assistance. We thank Jeff Sample for MUTU I cells. We thank Molly Schineller and Emma Wolinsky for technical assistance. We thank Yijie Ma for bioinformatic assistance. RNAseq data was deposited in NIH GEO Omnibus, GSE148910.

## Author contributions

Y.Z., R.G., C.J, and S.T. performed and analyzed the CRISPR and biochemical experiments. C.J and S.J.T. performed the RNAseq experiments, which were analyzed by R.G. and M.T. Y.N. and B.Z. provided assistance with 293 BAC experiments. Bioinformatic analysis was performed by R.G. and M.T. B.E.G supervised the study.

